# Structural Determinants of Catalytic Bias in an AMP-Forming Acetyl-CoA Synthetase from *Syntrophus aciditrophicus*

**DOI:** 10.64898/2026.07.06.736832

**Authors:** Selena Yaghoubi, David M. Dinh, Leonard M. Thomas, Neil Q. Wofford, Michael J. McInerney, Elizabeth A. Karr

**Author notes:** Department of Chemistry, University of California, Davis, Davis, CA 95616, USA.

## Abstract

Acetyl-coenzyme A (CoA) is a central metabolic intermediate that links carbon and energy metabolism across all domains of life. The interconversion of acetate and acetyl-CoA is carried out by three enzyme pathways: acetate kinase/phosphotransacetylase, ADP-forming acetyl-CoA synthetase, and AMP-forming acetyl-CoA synthetase (Acs). Acs enzymes serve critical physiological roles across diverse organisms by catalyzing a reversible two-step reaction forming acetyl-CoA and AMP from acetate and ATP. Isolated from the wastewater reclamation facility in Norman, Oklahoma, *Syntrophus aciditrophicus* strain SB (*Sa*) thermodynamically favors synthesizing acetate and ATP from acetyl-CoA and AMP using an AMP-forming acetyl-CoA synthetase (*Sa*Acs1). The origin of the preference for AMP formation and the structural determinants of both the thioester-forming step and catalytic bias remain poorly understood. Here, we report a 2.2 Å crystal structure of full-length *Sa*Acs1 in the adenylation conformation with acetyl-AMP bound in the active site. Structural comparison to the extensively characterized Acs enzymes from *Salmonella enterica* (*Se*Acs) and *Cryptococcus neoformans* (*Cn*Acs) revealed a displaced CoA-binding loop in *Sa*Acs1. Enzymatic assays support that *Sa*Acs1 preferentially catalyzes the ATP-forming reaction. Site-directed mutagenesis demonstrated that reversion of two residues, G196 and T197, at the beginning of the CoA-binding loop to the consensus sequence repositions the loop and shifts catalytic preference toward the AMP-forming direction. Together, these results establish the CoA-binding loop and G196 and T197 as the primary structural determinants of catalytic bias in *Sa*Acs1.

## Introduction

Acetyl-coenzyme A (CoA) is a central metabolic intermediate that links carbon and energy metabolism across all domains of life, serving as a key substrate in the tricarboxylic acid cycle, fatty acid biosynthesis, and a wide range of anabolic and catabolic pathways.^1, 2^ The reversible interconversion of acetate and acetyl-CoA is critical in allowing cells to activate acetate as a carbon source or to generate acetate and conserve energy depending on physiological demand (**Figure 1A**).^3^ Three enzymatic pathways carry out this reaction: acetate kinase/ phosphotransacetylase, ADP-forming acetyl-CoA synthetase, and AMP-forming acetyl-CoA synthetase (Acs) (**Figure 1B**).^4, 5^ Most fermentative bacteria rely on acetate kinase and phosphate acetyltransferase to synthesize ATP from acetyl-CoA, while acetate-producing archaea and certain eukaryotes use an ADP-forming acetyl-CoA synthetase.^6, 7^ The pathway a given organism employs is determined by its metabolic context, and in some cases, organisms have evolved to exploit these enzymes in ways that deviate from their canonical function.^6^

**Figure 1.**
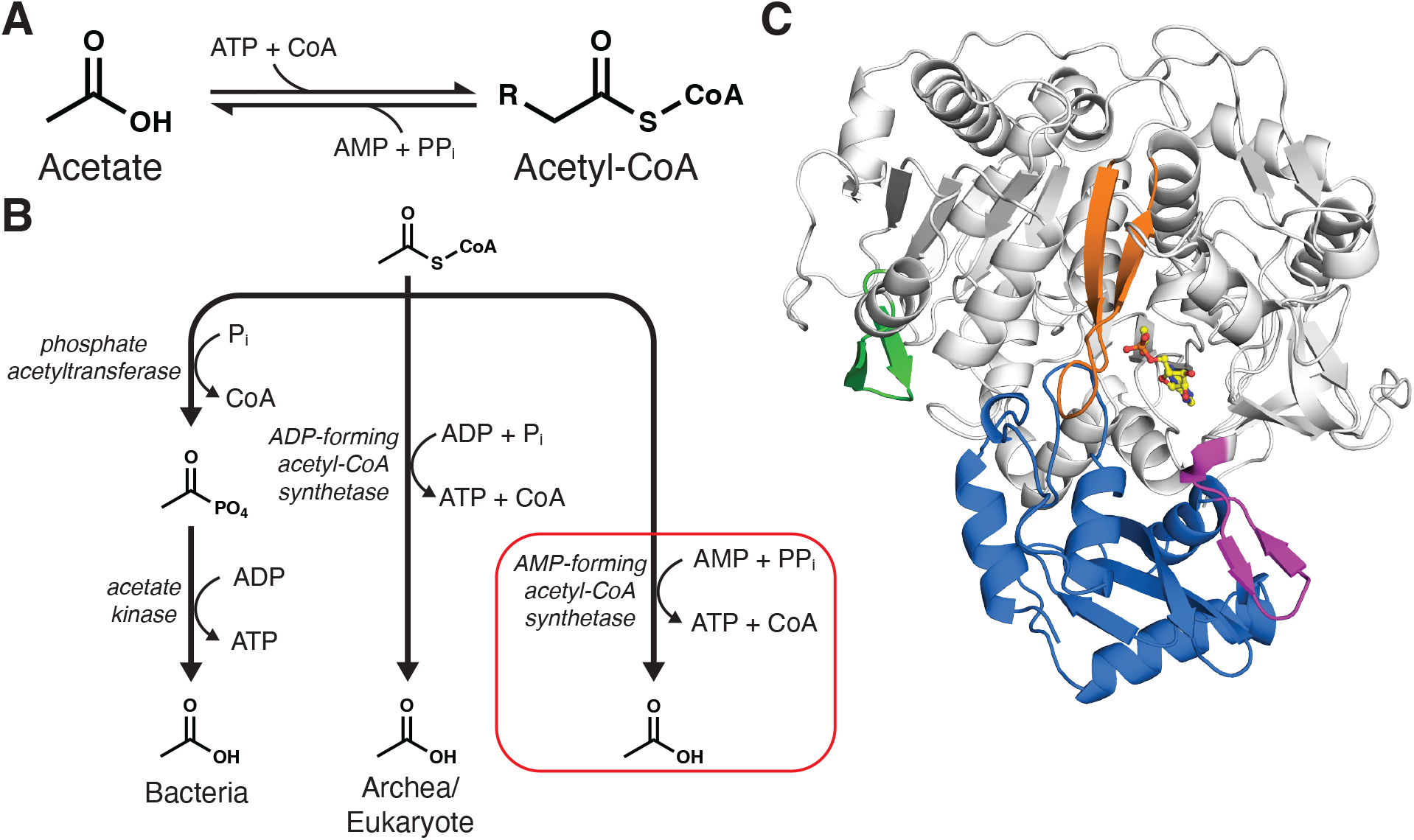
(**A**) The reversible reaction catalyzed by Acs enzymes. (**B**) Pathways of ATP synthesis by substrate-level phosphorylation with the pathway utilized by *S. aciditrophicus* boxed in red. (**C**) Full-length structure of *Sa*Acs1^WT^ with bound acetyl-AMP (yellow) resolved at 2.2 Å (PDB ID 9Y8G) with elements of secondary structure colored to represent the different regions of the protein. N-terminal domain (white), CoA binding loop (green), ATP binding loop (orange), hinge (magenta), and C-terminal domain (blue).

*Syntrophus aciditrophicus* strain SB (*Sa*) degrades aromatic and alicyclic compounds such as benzoate and cyclohexane carboxylate, respectively, in syntrophic association with hydrogen-consuming microorganisms (**Figure S1**).^8-10^ Because *Sa* lacks homologs of acetate kinase and phosphate acetyltransferase, it instead relies solely on an AMP-forming acetyl-CoA synthetase (*Sa*Acs1) to synthesize ATP from acetyl-CoA.^11, 12^ Although AMP-forming Acs enzymes typically catalyze the forward synthesis of acetyl-CoA and AMP from acetate and ATP, *Sa*Acs1 biases the reverse of this reaction, synthesizing acetate and ATP from acetyl-CoA and AMP. Prior characterization of *Sa*Acs1 demonstrated that the low ATP-to-AMP ratio and elevated pyrophosphate (PP^i^) levels in *Sa* favor operation of *Sa*Acs1 in the ATP-forming direction, with its activity sufficient to account for the observed rate of acetate production.^6, 13^ *Sa*Acs1 thus represents a unique case of evolutionary reprogramming within the Acs family. Rather than altering the underlying chemistry of the reversible Acs reaction, Acs enzymes have evolved distinct catalytic biases by redistributing catalytic efficiency between the ATP- and AMP-forming directions. The structural determinants responsible for tuning this catalytic bias, however, remain poorly understood.

Acs enzymes catalyze a reversible two-step reaction via an ordered Bi-Uni-Uni-Bi ping-pong mechanism.^2, 14^ In the first step, or adenylate-forming step, ATP binds the active site followed by acetate, resulting in formation of an enzyme-bound acetyl-AMP intermediate and release of pyrophosphate. This step proceeds through a single displacement mechanism in which acetate and pyrophosphate adopt an in-line, opposed orientation at the α-phosphate of ATP, with acetate acting as the nucleophile and pyrophosphate as the leaving group.^15, 16^ In the second step, or thioester-forming step, CoA binds the enzyme and acetyl-CoA is formed and released along with AMP (**Scheme 1 and Figures S2 and S3**).^17^ The mechanism by which Acs catalyzes this second half-reaction, however, remains poorly understood, and the structural features that govern the transition between the two half-reactions and determine the catalytic bias have not been fully established.^18^

**Scheme 1.**
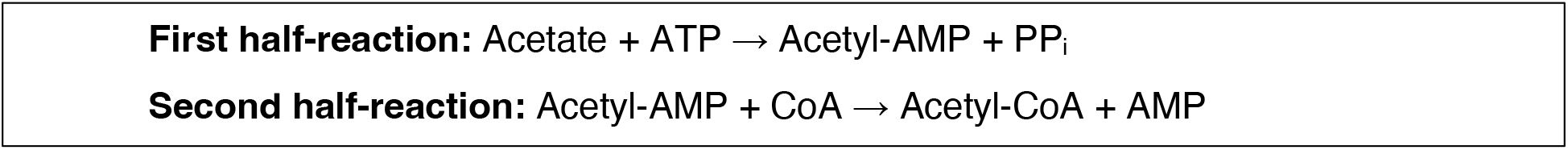
Half-reactions of acetate metabolism by Acs enzymes.

The adenylate-forming enzyme superfamily, of which *Sa*Acs1 is a member, shares ten regions of highly conserved residues (**Figure S4**).^19-21^ Structurally, Acs enzymes comprise a large N-terminal domain (fewer than 500 residues) and a smaller C-terminal domain (approximately 110 residues), with the active site at the domain interface (**Figure 1C**).^22-25^ The occupancy and identity of active site ligands influence the orientation of the C-terminal domain. A conserved residue within region 8 forms a hinge between the two domains and serves as a pivot point for an approximately 140° rotation of the C-terminal domain between the adenylation and thioesterification steps. ^26, 27^

To date, 24 Acs structures from nine organisms have been deposited in the PDB. Among these, a single apo structure (*Cn*Acs, PDB ID 5VPV), one ATP-bound structure (*Cn*Acs, PDB ID 5K8F), and one acetyl-AMP-bound structure (*Cn*Acs, PDB ID 7L4G) have been reported, while the remaining structures were co-crystallized with noncognate active site ligands (**Table S1**).^28^ Seven structures from *Cn*Acs and *Saccharomyces cerevisiae* Acs adopt the adenylate-forming conformation, while the remainder are in the thioester-forming conformation. Among the characterized Acs enzymes, *Salmonella enterica* Acs (*Se*Acs) and *Cn*Acs are the most extensively studied, with multiple structures determined for different ligand-bound states and detailed biochemical characterization of their substrate binding and conformational dynamics.^14, 18, 22, 28^ However, the limited number of structures capturing intermediate states of the reaction has left the structural basis of the thioester-forming step poorly understood. Furthermore, all previously determined Acs structures represent enzymes that have only been characterized in the canonical AMP-forming reaction, which raises the question of how an Acs enzyme functions in an organism that requires ATP production using an Acs enzyme.

Here, we report a structural and biochemical investigation of *Sa*Acs1, using *Se*Acs and *Cn*Acs as a comparative framework, aimed at identifying residues that were shown to contribute to catalysis in the ATP-forming direction. The crystal structure of wild-type *Sa*Acs1 was determined. Site-directed mutagenesis was used to introduce targeted substitutions at candidate residues, and the effects on enzyme structure and function were characterized. Structural and kinetic analyses together demonstrate that a loop adjacent to the CoA-binding pocket of *Sa*Acs1 is the primary structural determinant of its preference for the ATP-forming reaction.

## Materials and Methods

### Protein Expression and Purification

All constructs (*Sa*Acs1^WT^ and variants) were expressed in *E. coli* Rosetta™ (BL21 derivative, DE3) and induced at OD 0.6 with 0.1 mM IPTG overnight at 18°C. Cells were harvested by centrifugation, resuspended in equilibration buffer [50 mM Tris, 50 mM NaCl, 5 mM imidazole (pH 8)] with protease inhibitor tablets, and lysed by sonication. The soluble cell-free extract (CFE) was obtained by centrifugation and applied to a gravity column containing 1 mL Ni-NTA resin. The resin was washed sequentially with equilibration buffer, wash buffer 1 [50 mM Tris, 50 mM NaCl, 10 mM imidazole (pH 8)], and wash buffer 2 [50 mM Tris, 50 mM NaCl, 20 mM imidazole (pH 8)], and the protein was eluted stepwise with 5 mL of 100 mM imidazole in 50 mM Tris and 200 mM NaCl (pH 8), followed by 2 mL of 500 mM imidazole in 50 mM Tris and 300 mM NaCl (pH 8). Elution fractions were concentrated to approximately 500 μL and further purified by size exclusion chromatography (SEC) using a Superdex S-200 column equilibrated with 20 mM Tris and 150 mM NaCl (pH 8). The purified protein was concentrated to greater than 10 mg/mL prior to use (**Figure S5**).

### Crystallization of SaAcs1^WT^ and SaAcs1 Mutants

Purified recombinant *Sa*Acs1^WT^ (10 mg/mL) was incubated with 0.5 mM ATP and 0.5 mM acetate in microcentrifuge tubes for approximately one minute at room temperature. The microcentrifuge tube was dropped from benchtop height and centrifuged for 10 seconds, and an additional 0.5 mM ATP and 0.5 mM acetate was added just before tray setup. Initial thin needle-like crystals were obtained at room temperature by the sitting-drop method, mixing 1 μL of reservoir solution [0.1 M BisTris (pH 6.0), 0.2 M ammonium chloride, 22% (w/v) PEG 3350] with an equal volume of protein-substrate solution. To obtain larger crystals, initial crystals were transferred into fresh reservoir solution [0.1 M BisTris (pH 6.0), 22% (w/v) PEG 3350] for microseeding. The microseed stock was mixed with fresh protein-substrate solution and treated as above, with larger crystals obtained using reservoir solution [0.1 M BisTris (pH 6.0), 24% (w/v) PEG 3350, 0.2 M ammonium chloride]. *Sa*Acs1^WT^ crystals appeared within 24 hours of plating (**Figure S6**).

Variants *Sa*Acs1^G196E^, *Sa*Acs1^R199E^, *Sa*Acs1^K202E^, *Sa*Acs1^D527P^, and *Sa*Acs1^G196E/T197G^ (10 mg/mL) were crystallized using the same sitting-drop method and microseeding protocol as *Sa*Acs1^WT^, with incubation extended to approximately five minutes prior to tray setup. Crystals of *Sa*Acs1 were cryoprotected by serial soaking in reservoir solution with 5%, 10%, and 15% glycerol.

### Data Collections and Structure Determination

Diffraction data for the *Sa*Acs1^WT^, *Sa*Acs1^G196E^, *Sa*Acs1^K202E^, and *Sa*Acs1^D527P^ crystals were collected at the Stanford Synchrotron Radiation Lightsource (SLAC National Accelerator Laboratory) (Menlo Park, CA) at experimental station 9-2. Diffraction data for the *Sa*Acs1^G196E/T197G^ crystal was collected at the Biomolecular Structure Core (BSC – University of Oklahoma) (Norman, OK). The diffraction data for *Sa*Acs1^G196E/T197G^ was collected using a Rigaku MicroMax 007HF microfocus X-ray generator with an AFC11 4-axis goniometer and a Dectris Pilatus ® 200K silicon pixel detector. Data collection and processing were performed on Linux workstations using HKL3000R. The *Sa*Acs1 variant structures were determined using molecular replacement using the structure of *Sa*Acs1^WT^, where the wild-type structure was determined using a model predicted by AlphaFold 1.^29^ All data and refinement statistics are summarized in **Tables S2, S3, S4**.

### Protein Purification for Enzyme Assays and Kinetics

Proteins (*Sa*Acs1 WT and variants, *Se*Acs and variants, and *Cn*Acs) for enzyme assays and kinetics were purified by Ni-NTA affinity chromatography as described above, omitting the SEC step. Elution fractions were concentrated to approximately 500 μL and desalted into 50 mM Tris (pH 8) using 5 mL desalting columns, with protein elution confirmed by absorbance at 280 nm. Desalted protein samples were kept on ice until use and protein purity was assessed by SDS-PAGE.

### Enzyme Kinetics

Enzyme kinetics were determined in the AMP-forming and ATP-forming directions by varying one substrate within each reaction while holding the remaining two constant (**Figure S7**). All reactions were measured at a total volume of 400 μL and were preincubated in ultra-micro cuvettes at 37°C in a water bath. The reaction was initialized with the addition of purified enzyme in a buffer containing 50 mM Tris pH 8.5. Each reaction was monitored at 340 nm for 3 minutes using a Biochrom WPA Biowave II UV/visible spectrophotometer. The initial velocity data was then analyzed using GraphPad Prism to obtain apparent *K*_*M*_ (Michaelis-Menten constant) and *k*_*cat*_ (catalytic constant) of each substrate measured using non-linear regression.

In the AMP-forming direction, ATP kinetic constraints were measured by holding CoA and acetate constant at 480 μM and 5 mM, respectively, while ATP was varied from 25 to 5000 μM. CoA kinetic constraints were measured by holding both ATP and acetate constant at 5 mM, while CoA was varied from 12 to 600 μM. Acetate kinetic constraints were measured by holding CoA and ATP constant at 480 μM and 5 mM, respectively, while acetate was varied from 25 to 5000 μM. Each reaction contained 50 mM Tris-HCl buffer (pH 8.5) with 10 mM magnesium chloride, 1 mM phosphoenolpyruvate (PEP), 375 μM NADH, 2.8 U myokinase (MK), 2.2 U pyruvate kinase (PK), and 2.2 U lactate dehydrogenase (LDH).

In the ATP-forming direction, PP_i_ kinetic constraints were measured by holding AMP and acetyl-CoA constant at 2 mM and 0.5 mM, respectively, while PP_i_ was varied from 25 to 1250 μM. AMP kinetic constraints were measured by holding both PP_i_ and acetyl-CoA constant at 1 mM and 5 mM, respectively, while AMP was varied from 25 to 2500 μM. Acetyl-CoA kinetic constraints were measured by holding PP_i_ and AMP constant at 1 mM and 2 mM, respectively, while acetyl-CoA was varied from 25 to 500 μM. Each reaction contained 20 mM triethanolamine buffer (pH 7.8) with 3 mM magnesium chloride, 400 μM NADP+, 5 mM D-glucose, and 15 μg glucose-6-phosphate dehydrogenase/hexokinase (G6PDH/HK).

Because Acs enzymes follow an ordered Bi-Uni-Uni-Bi ping-pong mechanism, varying a single substrate while maintaining the remaining substrates at saturating concentrations yields *apparent* steady-state kinetic parameters (operational quantities obtained under saturating cosubstrate conditions) rather than microscopic rate constants. Throughout this work, *K*_*M*_, *k*_*cat*_, and *k*_*cat*_*/K*_*M*_ therefore refer to *apparent* steady-state parameters that provide a consistent basis for comparing catalytic performance and catalytic bias among Acs homologs and variants.^30, 31^

## Results

### Structural Comparison of SaAcs1 to SeAcs and CnAcs

*Sa*Acs1_WT_ (PDB ID 9Y8G) was crystallized in the presence of acetate and ATP. As a result, the structure exhibited electron density consistent with a molecule of acetyl-AMP bound within the active site (**Figure S8**). This step represents the end state of the first half-reaction for typical Acs enzymes and the initial state of the second half-reaction for *Sa*Acs1. Comparing the sequences of other Acs proteins to *Sa*Acs1, there are two conserved residues at the beginning of the loop that interact with the CoA molecule and are not conserved in *Sa*Acs1 (**Figure 2**).

**Figure 2.**
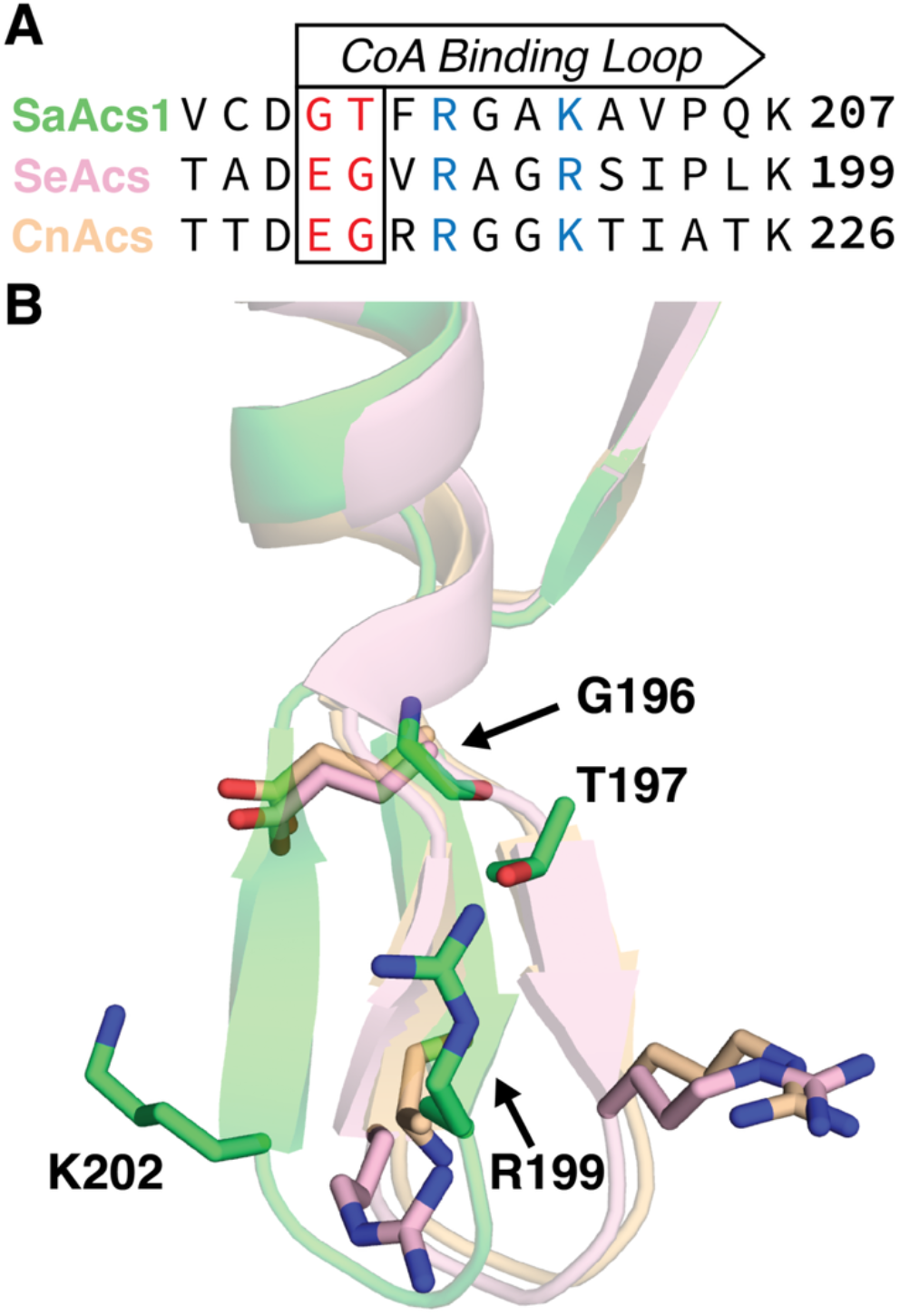
(**A**) Sequence alignment of the CoA-binding loop region for *Sa*Acs1, *Se*Acs, and *Cn*Acs with the residues at the beginning of the loop denoted in red and the conserved positively charged residues denoted in blue. (**B**) CoA-binding loop structural alignment of *Sa*Acs1^WT^ (green), *Se*Acs (salmon), and *Cn*Acs (tan). Critical residues at the beginning and within the loop are shown as sticks.

The structures of *Se*Acs and *Cn*Acs have been determined in the presence and absence of CoA, and reveal little to no change in the position of the CoA-binding loop during co-crystallization (**Figure S9**).^18, 28^ In contrast, one striking difference in the structure of *Sa*Acs1 was a shift in the position of the CoA loop (**Figure 2B**). While these two residues differ from those in *Se*Acs and *Cn*Acs, these positions are well conserved across other species (**Figure S10**). In addition to this shift, two well-conserved positively charged amino acids inside the loop known to interact with the nucleotide end of the CoA molecule are also displaced in *Sa*Acs1 compared to all other Acs structures. This difference in conformation of *Sa*Acs1^WT^ expands the entrance to the CoA/acetyl-CoA binding pocket compared to *Se*Acs and *Cn*Acs, which exhibit more restricted access to their CoA binding cleft (**Figure S11**).

To test the proposal that the structural shift of the loop was due to the presence of G196 and T197, two variants were prepared that reverted these residues to the sequences of *Se*Acs and *Cn*Acs, *Sa*Acs1^G196E^ and *Sa*Acs1^G196E/T197G^. While structures of both variants were determined, *Sa*Acs1^G196E^ (PDB ID 36UO) showed no significant difference in the position of the CoA binding loop when aligned to *Sa*Acs1^WT^ (**Figure S12A**). However, *Sa*Acs1^G196E/T197G^ (PDB ID 36UT) revealed a 2.1 Å shift of the loop compared to *Sa*Acs1^WT^, towards the positions observed in *Se*Acs and *Cn*Acs in support of the proposal that the two residues drive the differences between structures (**Figure 3A**). Further support that the double mutation is necessary for the conformational shift comes from the reorientation of the two positively charged amino acids within the loop. In *Sa*Acs1^G196E^, residues 199 and 202 remained in the same position observed in *Sa*Acs1^WT^, while the same residues of *Sa*Acs1^G196E/T197G^ display the same orientation as R191 and R194 of *Se*Acs and R218 and K221 of *Cn*Acs, respectively (**Figure 3B**). Additionally, structures of *Sa*Acs1^K202E^ (PDB ID 36UR), and *Sa*Acs1^D527P^ (PDB ID 36US) were resolved and showed no difference in the position of the CoA binding loop when aligned to *Sa*Acs1^WT^, indicating these residues do not induce a conformational shift (**Figures S12B and C**).

**Figure 3.**
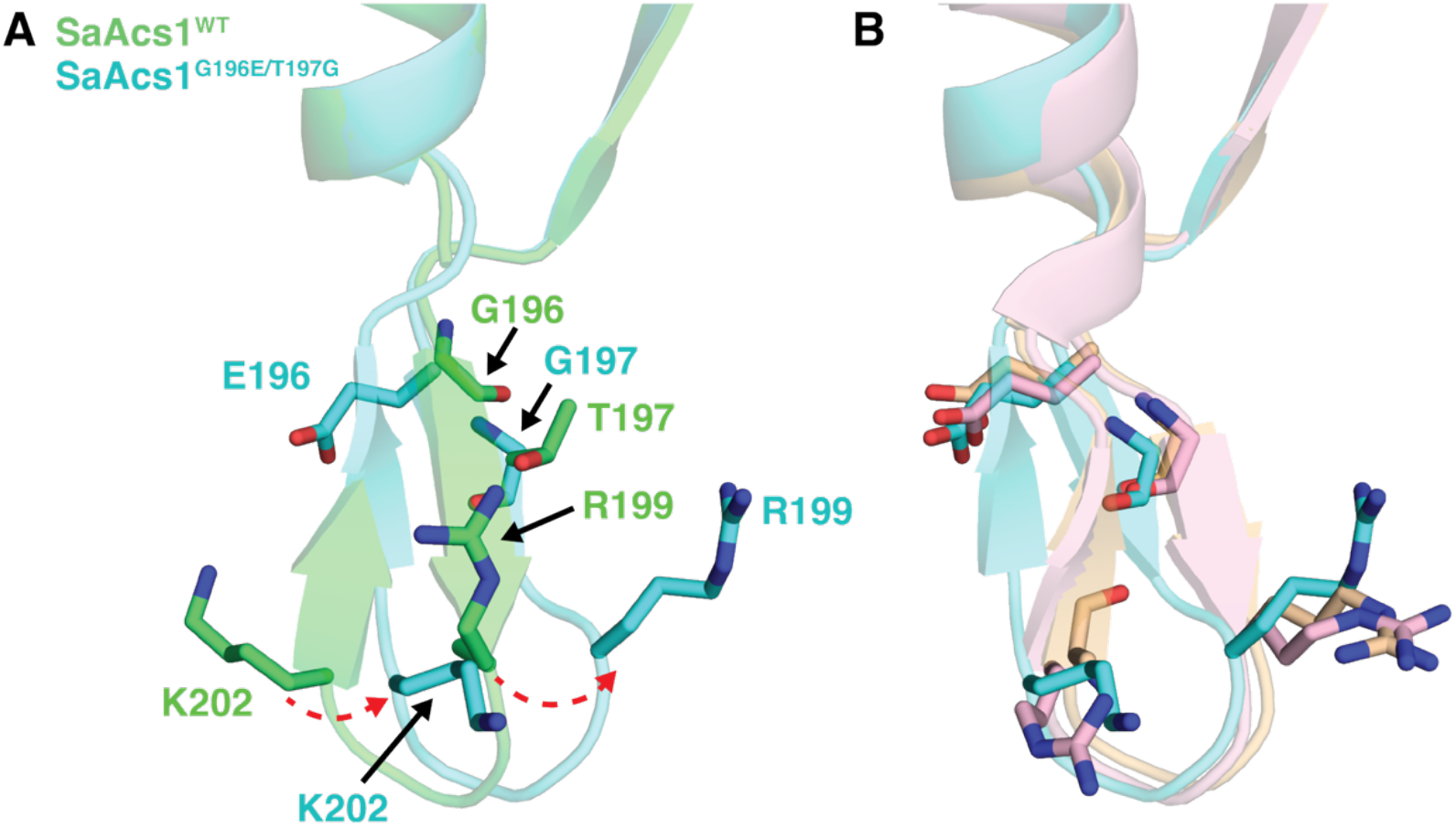
(**A**) CoA-binding loop alignment of *Sa*Acs1^WT^ (green) and *Sa*Acs1^G196E/T197G^ (cyan). Movement of residues R199 and K202 from WT to G196E/T197G are depicted in red dashed arrows. (**B**) CoA-binding loop alignment of *Sa*Acs1^G196E/T197G^ (cyan) to *Se*Acs (salmon) and *Cn*Acs (tan). Critical residues at the beginning and within the loop are shown as sticks.

### Enzyme Kinetics

To understand how the structure of *Sa*Acs1 preferentially promotes the formation of acetate and ATP over the production acetyl-CoA and AMP, typical of other Acs enzymes, the activity of *Sa*Acs1^WT^ was assessed against *Se*Acs and *Cn*Acs. Apparent steady-state kinetic parameters (*K*_*M*_, *k*_cat_, and *k*_cat_/*K*_*M*_) were determined for each substrate in both reaction directions and used to compare catalytic efficiencies among Acs homologs and variants.^14, 18^ The validity of Michaelis-Menten parameters for a reversible ordered Bi-Uni-Uni-Bi ping-pong mechanism must be treated with caution due their sensitivity to assay conditions including substrate concentration (see Supplemental Information for details).^30, 31^ The value of these apparent kinetic parameters is therefore to serve as a qualitative marker for catalytic bias, allowing relative ratios between forward and reverse reactions to be compared between variants. In addition, they enable comparisons to be made across the acyl- and aryl-CoA subfamily, where determination of these apparent values are performed as a standard assay.^18, 32-35^

The three substrates involved with the forward (AMP-forming) reaction include acetate, ATP, and CoA. The three substrates involved with the reverse (ATP-forming) reaction include pyrophosphate (PP_i_), AMP, and acetyl-CoA. *Sa*Acs1 displayed a greater apparent *K*_M_ for ATP than for AMP, while both *Se*Acs and *Cn*Acs had higher apparent *K*_M_ for AMP compared to that for ATP, indicating their difference in preference for the reverse versus forward reactions, respectively. However, all three Acs enzymes showed similar apparent *k*_*cat*_ values for the six substrates, with *Sa*Acs1 having a slightly greater *k*_*cat*_ for ATP and *Cn*Acs exhibiting a greater *k*_*cat*_ for CoA. Overall, the apparent catalytic efficiency for AMP was greater than ATP for *Sa*Acs1, while the efficiency for ATP was greater than AMP for *Se*Acs and *Cn*Acs, where differences between the enzymes were primarily driven by the apparent *K*_*M*_.

To assess whether the residues at the beginning of the CoA binding loop could drive the catalytic bias toward the AMP-forming direction as well as the positioning of the structural motif, five variants were prepared for *Sa*Acs1 and *Se*Acs: *Sa*Acs1^G196E^, *Sa*Acs1^G196E/T197G^, *Sa*Acs1^G196E/T197G/K202R^, *Se*Acs^E188G/G189T^, and *Se*Acs^E188G/G189T/R194K^. The activity of these variants was compared to the WT Acs enzymes (**Figure 4 and Table S5**).

**Figure 4.**
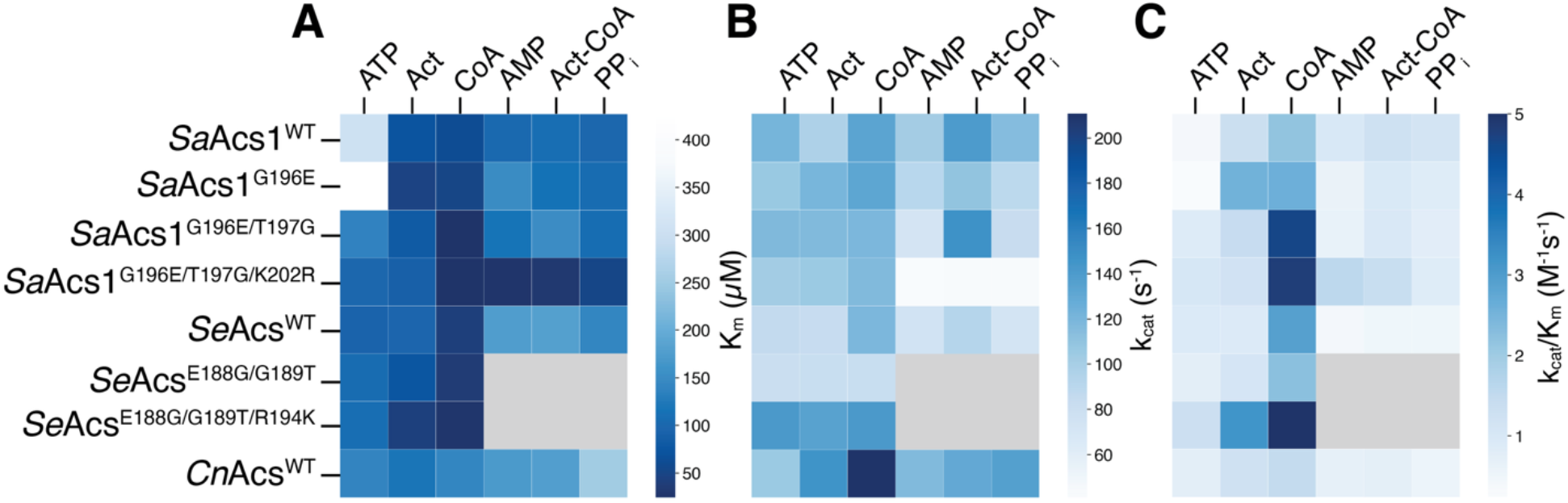
Heatmap analysis of apparent Michaelis-Menten parameters for *Sa*Acs1^WT^, *Sa*Acs1 CoA-binding loop variants, *Se*Acs^WT^, and *Cn*Acs^WT^. Apparent (**A**) *K*_*M*_ (μM), (**B**) *k*_*cat*_ (s^-1^), and (**C**) *k*_*cat*_/*K*_*M*_ (M^-1^s^-1^) values are shown for all six substrates across enzymes. In each map, the left three columns (ATP, acetate (Act), CoA) represent AMP-forming substrates and the right three columns (AMP, acetyl-CoA (Act-CoA), PP_i_) represent ATP-forming substrates. Gray cells indicate no detectable activity during the time course of the measurement. Color gradient represents the magnitude of each parameter.

In the AMP-forming reaction, both *Sa*Acs1 variants displayed apparent greater catalytic efficiency for ATP (G196E/T197G: 2.15-fold and G196E/T197G/K202R: 2.58-fold) and CoA (G196E/T197G: 2.14-fold and G196E/T197G/K202R: 2.23-fold) compared to *Sa*Acs1^WT^. For *Se*Acs^E188G/G189T/R194K^, greater apparent catalytic efficiency was observed for all three substrates in the AMP-forming reaction compared to *Se*Acs^WT^. In the ATP-forming reaction, the catalytic efficiencies for all substrates with *Sa*Acs1^G196E^ and *Sa*Acs1^G196E/T197G^ resembled *Sa*Acs1^WT^. Conversely, *Sa*Acs1^G196E/T197G/K202R^ displayed moderately greater apparent efficiencies for AMP (1.23-fold) and acetyl-CoA (1.42-fold) compared to *Sa*Acs1^WT^. For the *Se*Acs variants there was no detectable NADPH formation in the time course of the reaction.

While the positioning of the loop and the activity bias was clearly dominated by the residues at the beginning of the loop, the function of other potential key amino acid residues within the CoA-binding loop was also investigated in the AMP-forming and ATP-forming directions. The apparent catalytic efficiencies for all variants within the CoA-binding loop (*Sa*Acs1^G196E^, *Sa*Acs1^G196E/T197G^, *Sa*Acs1^K202A^, *Sa*Acs1^K202R^, *Sa*Acs1^K202E^, *Sa*Acs1^K202S^, *Sa*Acs1^K202Q^,*Sa*Acs1^R199A^, *Sa*Acs1^R199E^) were compared to *Sa*Acs1^WT^. In the ATP-forming direction, *Sa*Acs1^K202A^, *Sa*Acs1^K202E^, *Sa*Acs1^R199A^, and *Sa*Acs1^R199E^ exibited no detectable NADPH formation in the time course of the reaction (**Figure 5 and Table S6**).

**Figure 5.**
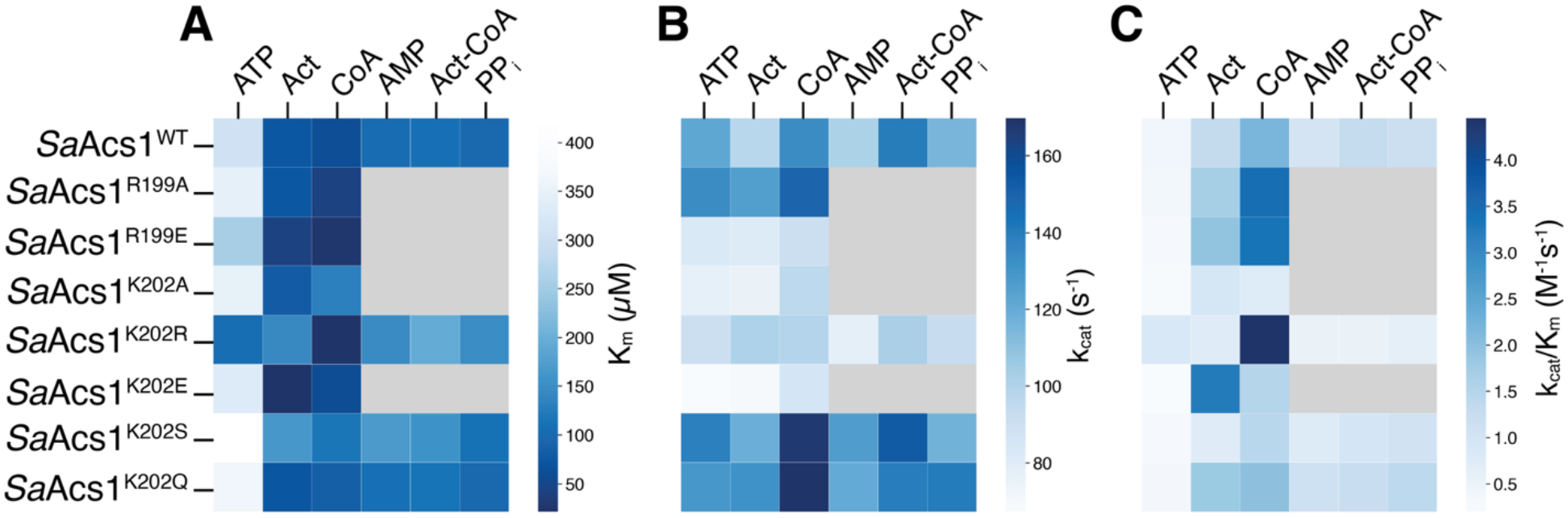
Heatmap analysis for *Sa*Acs1^WT^ and CoA-binding loop variants at R199 and K202. Apparent (**A**) *K*_M_ (μM), (**B**) *k*_cat_ (s^-1^), and (**C**) *k*_*cat*_/*K*_M_ (M^-1^s^-1^) values are shown for all six substrates across enzyme variants. In each map, the left three columns (ATP, acetate (Act), CoA) represent AMP-forming substrates and the right three columns (AMP, acetyl-CoA (Act-CoA), PP_i_) represent ATP-forming substrates. Gray cells indicate no detectable activity during the time course of the measurement. Color gradient represents the magnitude of each parameter.

## Discussion

The interconversion of acetate and acetyl-CoA is essential to carbon and energy metabolism across diverse organisms. Three enzyme pathways carry out this reaction: acetate kinase/phosphotransacetylase, ADP-forming acetyl-CoA synthetase, and AMP-forming acetyl-CoA synthetase (Acs).^7, 36^ Because *S. aciditrophicus* lacks homologs of acetate kinase and phosphate acetyltransferase, it relies solely on *Sa*Acs1 to produce ATP from acetyl-CoA, AMP, and pyrophosphate.^6^ While AMP-forming Acs enzymes typically synthesize acetyl-CoA and AMP from acetate and ATP, *Sa*Acs1 must support the reverse reaction based on *in vivo* concentration measurements.^6^ Using X-ray crystallography and site-directed mutagenesis of *Sa*Acs1 and its homologs, this study supports the argument that this activity is driven by evolutionary sequence divergences of *Sa*Acs1 rather than purely environmental substrate concentrations. Specifically, the positioning of CoA-binding loop was identified as a key structural motif that reconciles this shift.

Sequence alignment of *Sa*Acs1 with *Se*Acs, *Cn*Acs, and *Sc*Acs revealed approximately 50% sequence identity across comparisons. Two highly conserved residues at the beginning of the CoA-binding loop, a glutamic acid and glycine in most Acs enzymes, are replaced by G196 and T197 in *Sa*Acs1. Although this loop region falls outside the ten conserved adenylate-forming motifs and is not conserved within the acyl- and aryl-CoA subfamily, its positioning influences catalysis. While G196 and T197 are not structurally shifted in *Sa*Acs1, the conserved charged residues within the loop, R199 and K202, are displaced from the positions of their corresponding residues in *Se*Acs (R191, R194) and *Cn*Acs (R218, K221). Given that these residues have an established role in CoA binding and stabilization of the thioester-forming conformation, this displacement in *Sa*Acs1 likely weakens CoA interactions and expands the CoA-binding pocket. This expansion may facilitate the entry of the acetyl-CoA substrate in the ATP-forming direction, which is supported, in part, by *Sa*Acs1^WT^ displaying greater apparent *K*_*M*_ and lower catalytic efficiency for AMP relative to ATP, while *Se*Acs and *Cn*Acs showed the opposite.

Determination of the structure of the double variant *Sa*Acs1^G196E/T197G^ revealed a repositioned CoA loop where R199 and K202 were restored to orientations which structurally align with the corresponding residues in *Se*Acs and *Cn*Acs. However, the single variant, *Sa*Acs1^G196E^, did not exhibit a substantial deviation from WT, confirming that both residues are necessary to drive the conformational shift. Kinetically, *Sa*Acs1^G196E/T197G^ showed increased apparent catalytic efficiencies for all AMP-forming substrates and decreased efficiencies in the ATP-forming direction, resembling *Se*Acs^WT^ and *Cn*Acs^WT^. As mentioned above, Michaelis-Menten steady state parameters offer limited quantitative value in their microscopic interpretiblity for a reversible multistep mechanism like that of Acs enzymes due to their complexity and sensitivity to environmental conditions. However, determination of the apparent parameters do enable the calculation of an effective catalytic bias parameter, β, which is a product of the apparent catalytic efficiencies determined for each reaction component (**Eq. 1**).

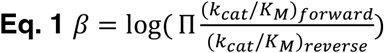

This parameter enables a simple qualitative comparison of the effective catalytic bias of each Acs enzyme, where β > 1 is biased towards AMP-formation and β < 1 is biased towards ATP-formation. Errors associated with each β-value were propogated from the corresponding standard errors of the respective kinetic measurements used for its calculation. Calculation of β for the variants prepared in this study demonstrate that *Cn*Acs and *Se*Acs exhibit strong biases towards AMP-formation with β = +0.78 and β = +1.40, respectively, while *Sa*Acs1 exhibits limited bias with a lean in favor of ATP-formation (β = -0.08) (**Figure 6**). The utility of this analysis is clear upon comparison with the double and triple variants, *Sa*Acs1^G196E/T197G^ and *Sa*Acs1^G196E/T197G/K202R^, where β increases significantly to +1.10 and +0.52, respectively, revealing a bias toward AMP-formation consistent with the structural repositioning of the CoA-binding loop. While the *Sa*Acs1^G196E^ and *Sa*Acs1^K202R^ variants did not show structural deviations of their CoA loops, their activity demonstrated a shift in functional bias towards AMP-formation.

**Figure 6.**
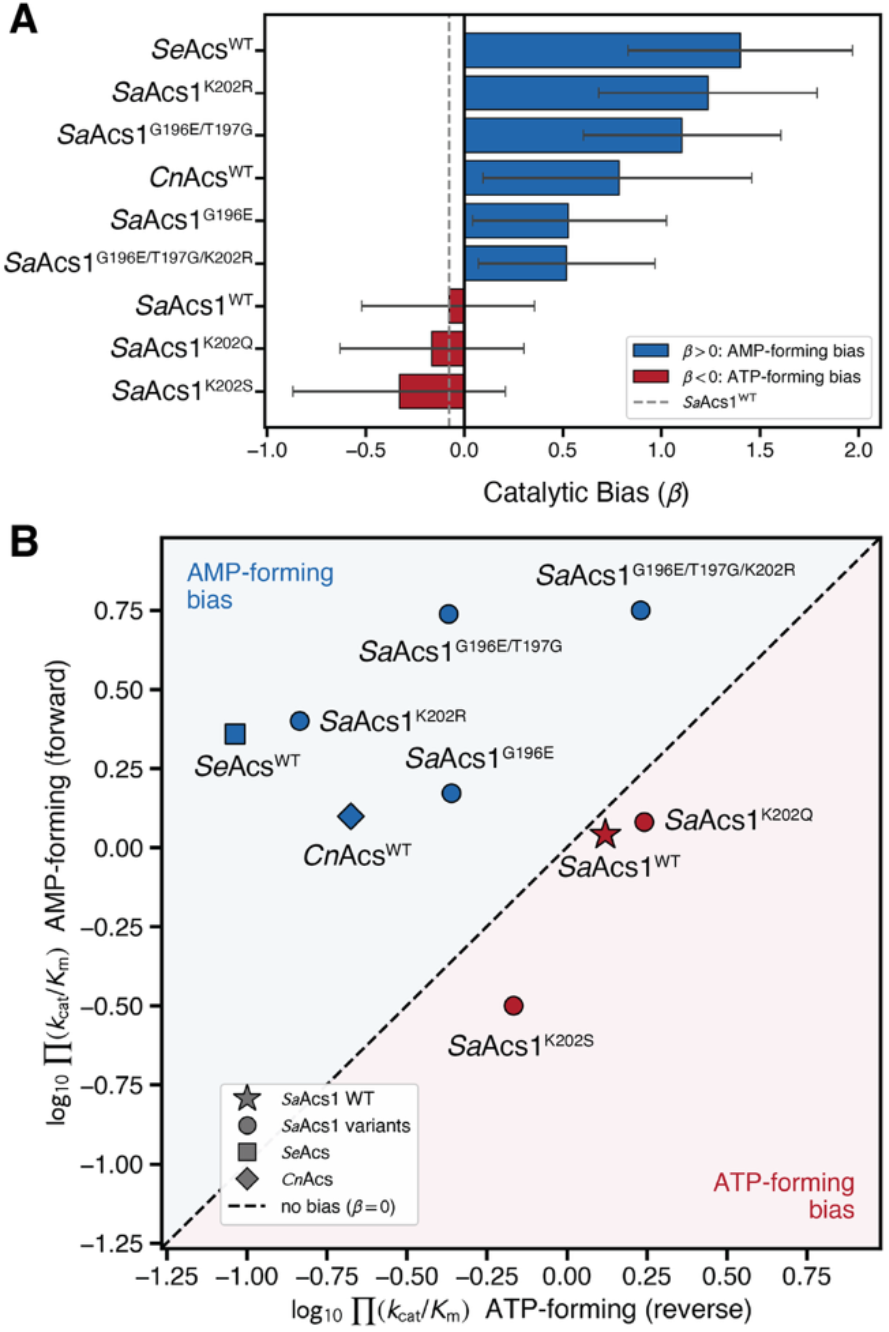
(**A**) Bar graph plot depicting effective catalytic bias, β, of variants where apparent steady state parameters were measurable. Error bars represent propagated error from standard errors of the kinetic experiments. (**B**) Bias comparison plot of overall forward and reverse apparent *k*_*cat*_/*K*_*M*_ contributions to β. *Sa*Acs1^WT^ is depicted as a star with variants depicted as circles, *Se*Acs^WT^ as a square, *Cn*Acs as a diamond.

Further mutagenesis of R199 and K202 confirmed their importance within the loop. Variants *Sa*Acs1^K202A^, *Sa*Acs1^K202E^, *Sa*Acs1^R199A^, and *Sa*Acs1^R199E^ showed no detectable ATP-forming activity within the time-frame of the experiment, preventing the calculation of β. It is interesting to note that ATP-formation was sufficiently impacted by charge-reversal and alanine mutations of R199 and K202, while swapping positively-charged residues exhibited limited changes. For example, *Sa*Acs1^K202R^ retained ATP-forming activity, and calculation of β revealed a bias toward AMP-formation (β = +1.24), indicating that the charge at position 202 influences the bias of the enzyme. Taken together, the structural and kinetic data establish the CoA-binding loop as a primary determinant of catalytic bias in *Sa*Acs1, with residues G196 and T197 modulating the positioning of R199 and K202 and their capacity for CoA interaction.

## Conclusions

The functional analysis of the residues within the CoA-binding loop supports the hypothesis that the structural changes observed for *Sa*Acs1 affect the overall function by redistributing their catalytic efficiency in favor of the ATP-forming reaction, relative to other Acs enzymes. The structure of *Sa*Acs1^WT^ revealed that there is a larger CoA-binding pocket which would allow enhanced entry and affinity for acetyl-CoA. *Sa*Acs1^G196E/T197G^ demonstrated that changing the residues at the beginning of the CoA loop shifted its position to align with *Se*Acs and *Cn*Acs and switched the functional bias of the enzyme to operating in the AMP-forming direction, similar to other Acs enzymes. Additionally, the biochemical analysis of the conserved positive residues, R199 and K202, within the CoA loop confirmed that their shift in the structure of *Sa*Acs1 impacts the interactions of the residues and likely plays an important role in binding substrates in the CoA-binding pocket.

## Supporting Information

Additional data, figures, crystallographic data collection, and enzyme kinetics.

## Accession Codes

The crystal structures of *Sa*Acs1^WT^ (PDB ID: 9Y8G), *Sa*Acs1^G196E^ (PDB ID: 36UO), *Sa*Acs1^G196E/T197G^ (PDB ID: 36UT), *Sa*Acs1^K202E^ (PDB ID: 36UR), and *Sa*Acs1^D527P^ (PDB ID: 36US)were deposited in the PDB and can be accessed free of charge via https://www.rcsb.org/. UniProt accession codes for *Sa*Acs1, *Se*Acs, and *Cn*Acs are Q2LWR8, Q8ZKF6, and J9VFT1, respectively.

## Notes

The authors declare no competing financial interest.

## Acknowledgements

The authors would like to acknowledge Prof Alec H. Follmer for his assistance in the preparation and assembly of this work. This work was supported by DOE Basic Energy Sciences Physical Biosciences Program Award DE-FG02-96ER20214 to Dr Elizabeth A. Karr, Professor and Vice Provost, and Dr Michael J. McInerney, Professor Emeritus, University of Oklahoma. The OU Biomolecular Sciences Core (BSC) was used for crystallization and data collection, and OU Protein Production and Characterization Core (PPCC) was used for protein purification. NIH COBRE Grant P20GM103640 and P30GM145423 funds both the OU BSC and PPCC. Use of the Stanford Synchrotron Radiation Lightsource, SLAC National Accelerator Laboratory, is supported by the U.S. Department of Energy, Office of Science, Office of Basic Energy Sciences under Contract No. DE-AC02-76SF00515. The SSRL Structural Molecular Biology Program is supported by the DOE Office of Biological and Environmental Research, and by the National Institutes of Health, National Institute of General Medical Sciences (P30GM133894).

