## Supplemental Information for "Structural Determinants of Catalytic Bias in an AMP-Forming Acetyl-CoA Synthetase from *Syntrophus aciditrophicus*"

### Table of Contents

Figure S1. Metabolism of *S. aciditrophicus*....S2
Figure S2. Synthesis of acetyl-CoA by Acs enzymes....S3
Figure S3. Enzymatic mechanism of Acs enzymes....S4
Figure S4. Multiple Sequence Alignment of Acs family proteins....S5
Table S1. List of all Acs proteins deposited in the Protein Data Bank....S8
Figure S5. *Sa*Acs1 protein purification....S9
Figure S6. *Sa*Acs1$^{WT}$ crystals....S10
Table S2. Crystallographic Data Collection and Refinement Statistics....S11
Table S3. Crystallographic Data Collection and Refinement Statistics....S12
Table S4. Crystallographic Data Collection and Refinement Statistics....S13
Figure S7. *Sa*Acs1 enzymatic activities for activity and kinetics assays....S14
Figure S8. Active site of *Sa*Acs1$^{WT}$....S15
Figure S9. CoA loop alignment of *Se*Acs and *Cn*Acs....S16
Figure S10. Acs CoA-binding loop alignment....S17
Figure S11. Comparison of the CoA-binding pocket surface....S18
Figure S12. Comparison of *Sa*Acs1 variants CoA loops....S19
Discussion of kinetic parameter determination....S20
Table S5. *Sa*Acs1$^{WT}$, *Sa*Acs1 CoA-binding loop variants, *Se*Acs$^{WT}$, and *Cn*Acs$^{WT}$ kinetics....S22
Table S6. *Sa*Acs1$^{WT}$ kinetics compared to *Sa*Acs1 CoA loop variants....S23

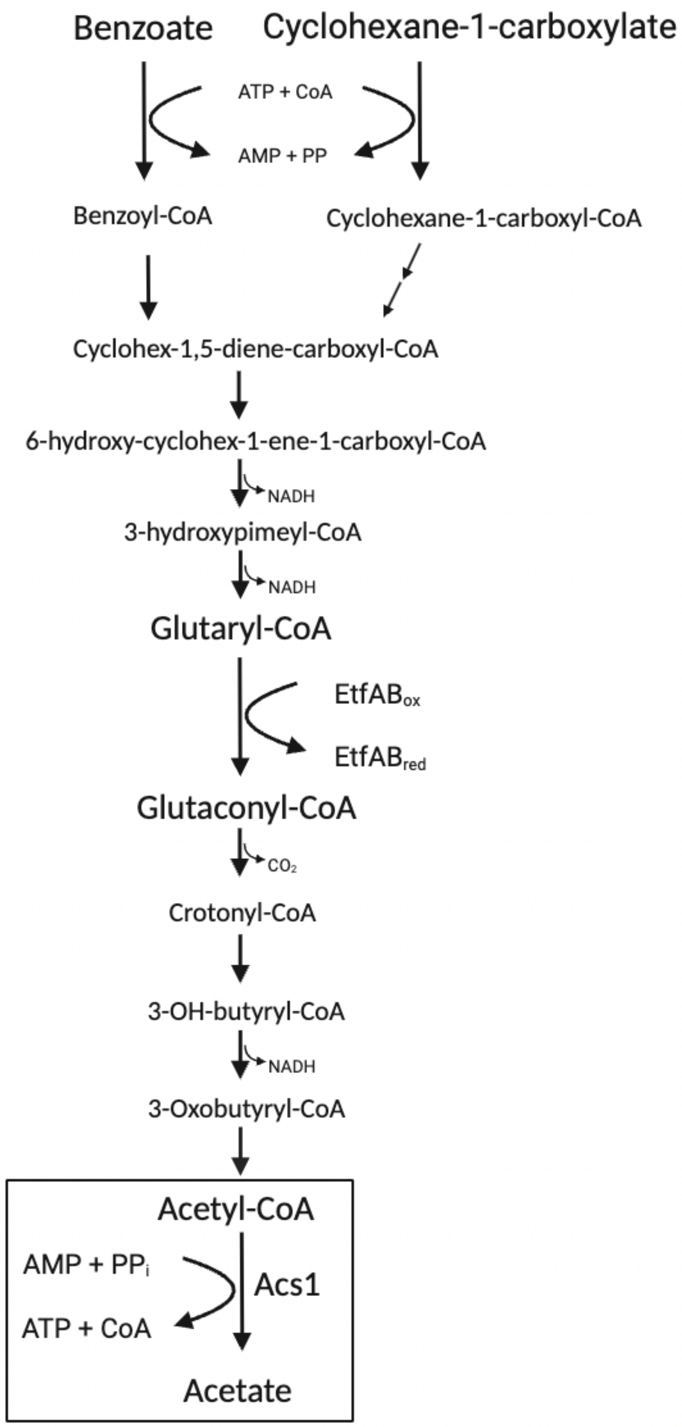


**Figure S1.** Metabolism of benzoate, cyclohexane-1-carboxylate, and crotonate by *S. aciditrophicus*.

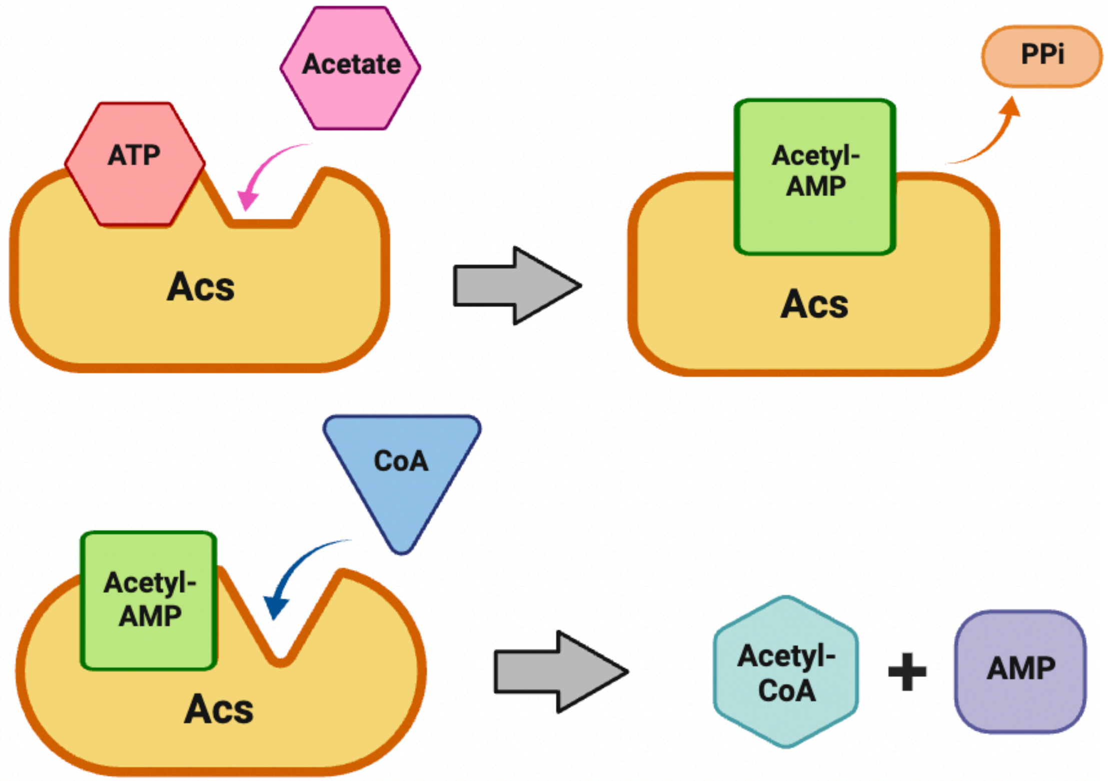


**Figure S2.** Synthesis of acetyl-CoA by AMP-forming acetyl-CoA synthetase. First half-reaction for the acetate activation to acetyl-AMP. Second half-reaction for the conversion of acetyl-AMP to acetyl-CoA.

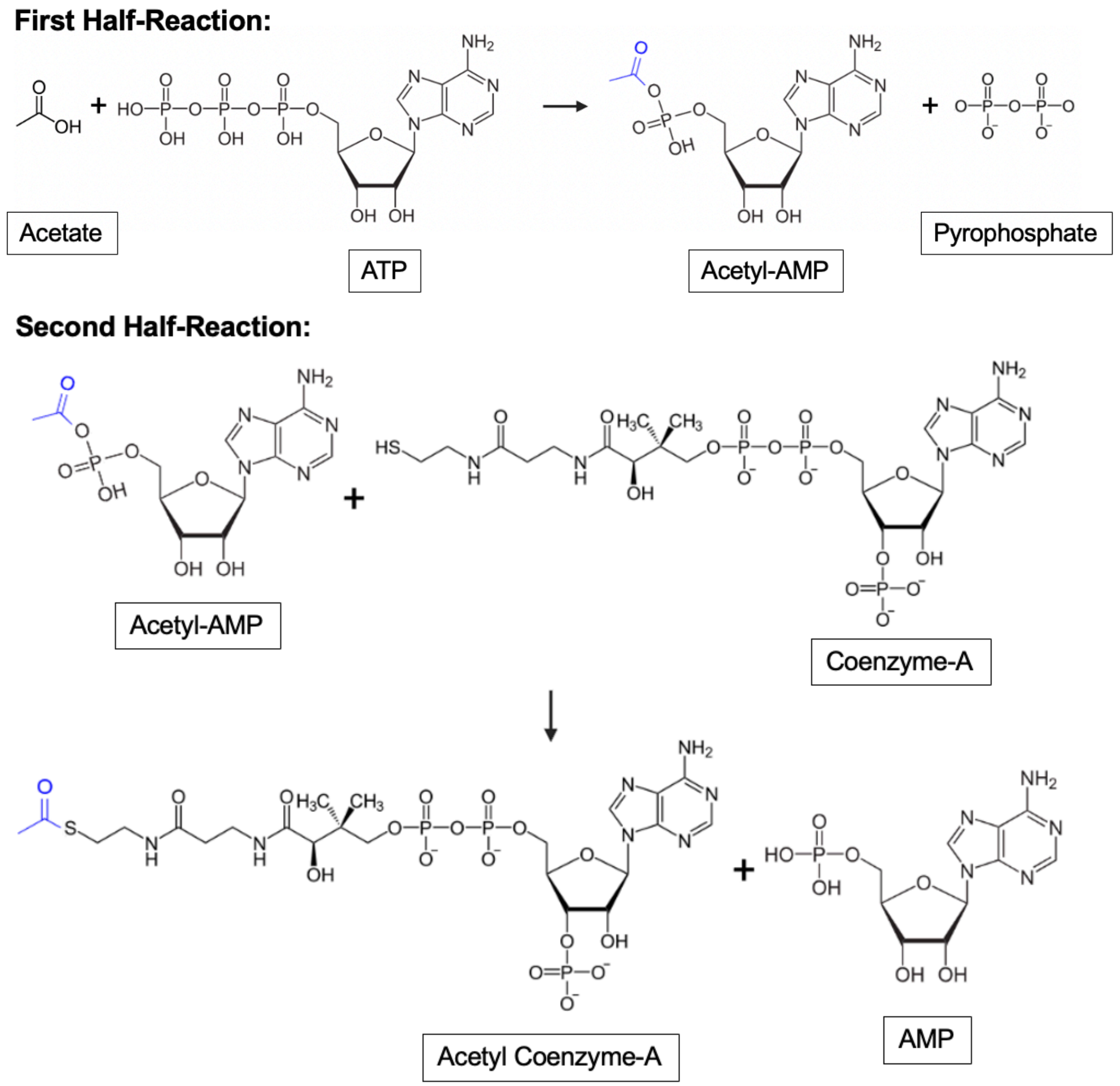


**Figure S3.** The reversible two-step enzymatic mechanism of AMP-forming acetyl-CoA synthetase. In the first half-reaction, an acetyl-adenylate intermediate (acetyl-AMP) is formed with the release of pyrophosphate. In the second half-reaction, acetyl-AMP intermediate then reacts with Coenzyme-A to form acetyl-CoA which then releases AMP.

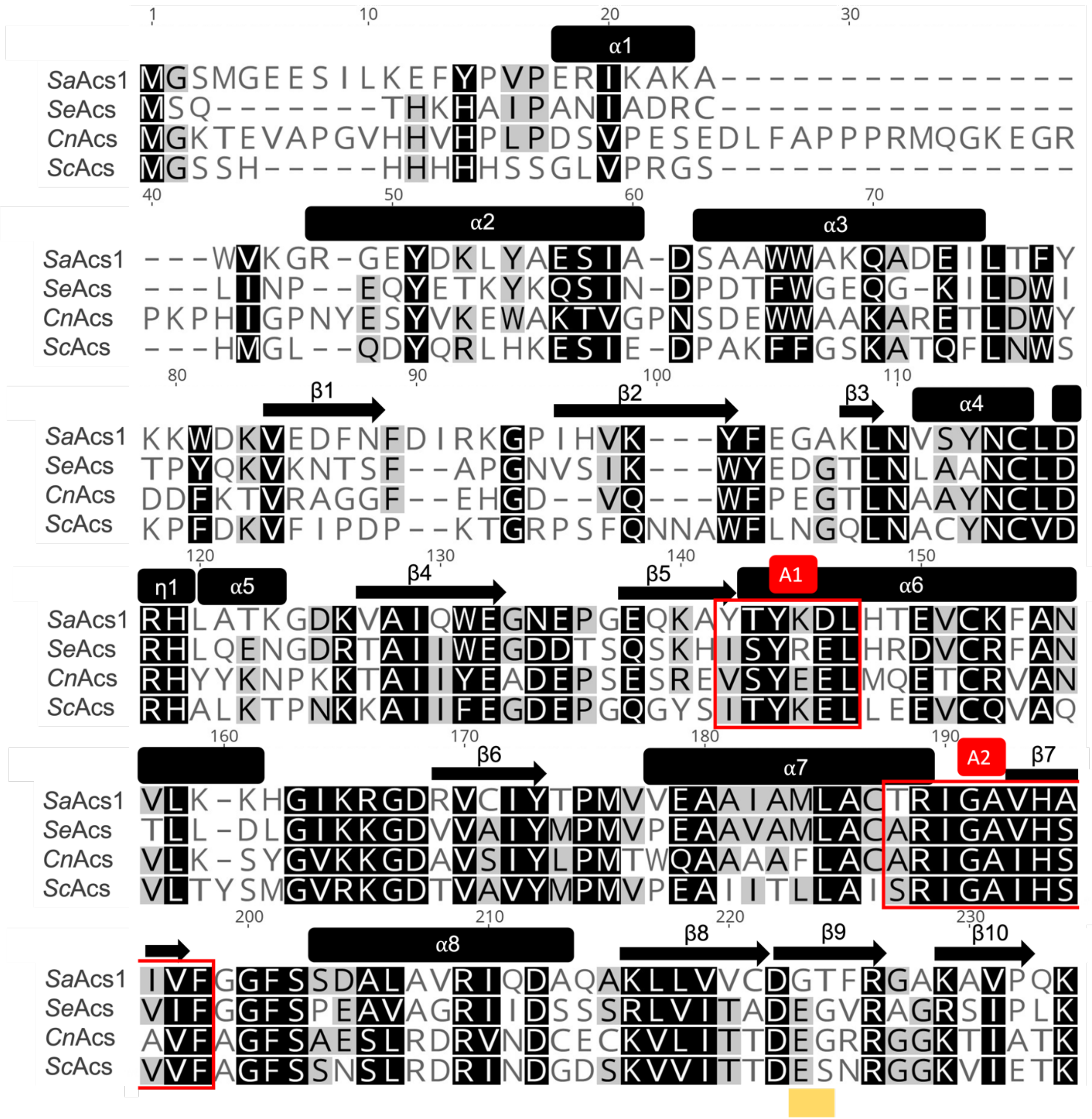

SaAcs1
SeAcs
CnAcs
ScAcs
α1
α2
α3
α4
α5
α6
α7
α8
η1
β1
β2
β3
β4
β5
β6
β7
β8
β9
β10
A1
A2

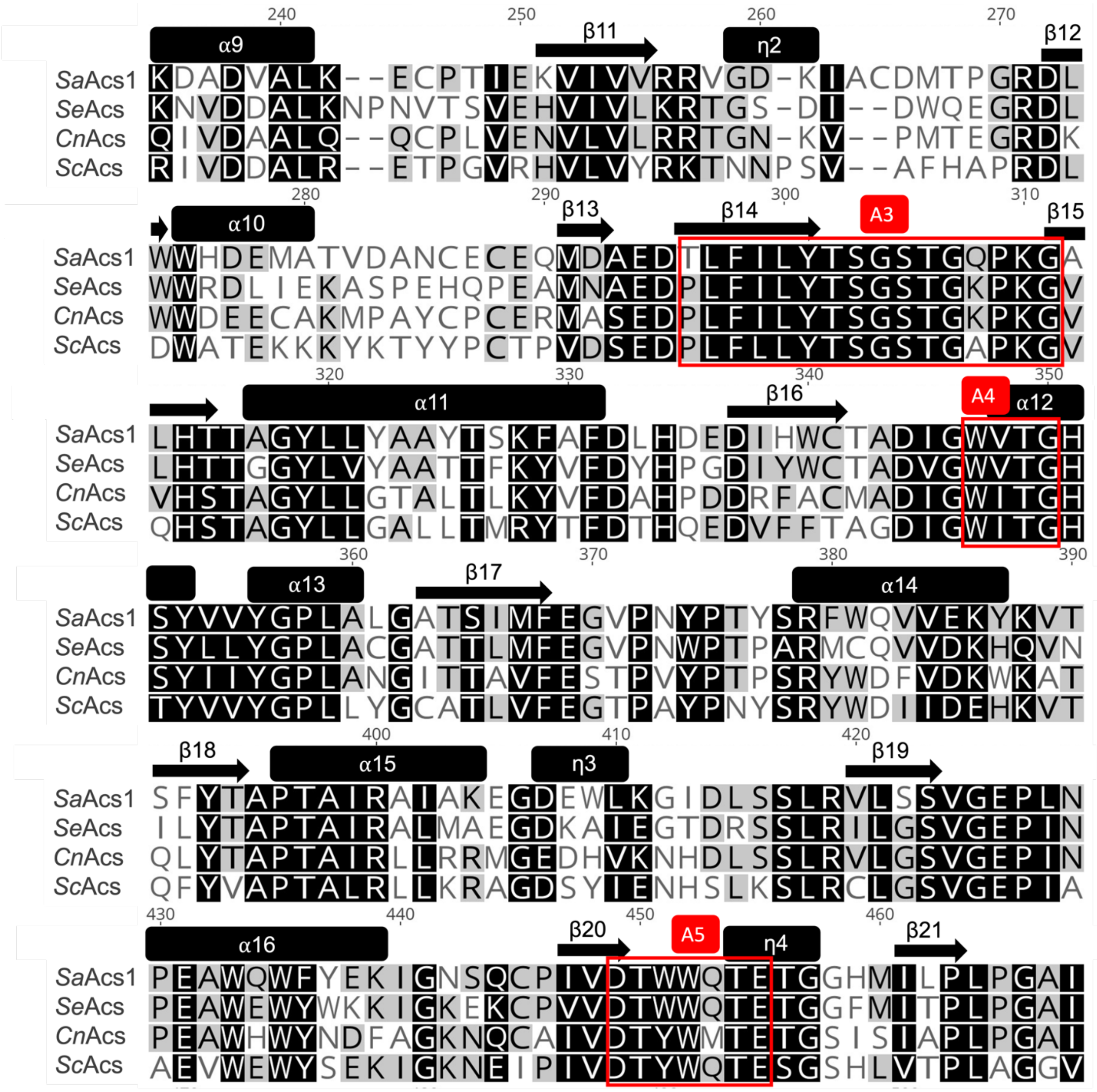

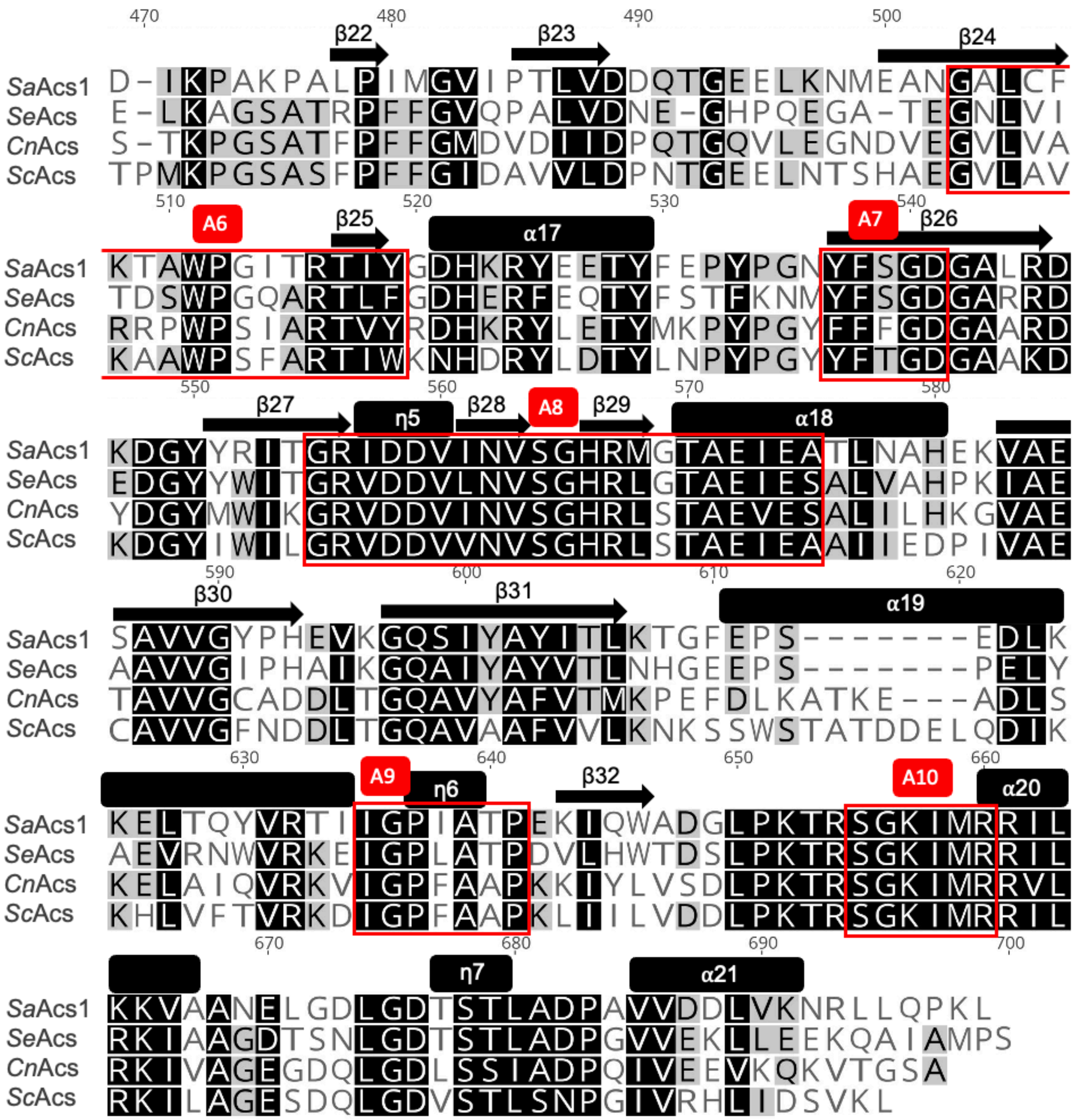

**Figure S4.** Multiple Sequence Alignment of Acs family proteins. Multiple sequence alignment by MAFFT of *Sa*Acs1, *Se*Acs (*Salmonella enterica*, PDB ID: 2P2F, sequence identity: 53.00%), *Cn*Acs (*Cryptococcus neoformans*, PDB ID: 5K8F, sequence identity: 47.38%), and *Sc*Acs (*Saccharomyces cerevisiae*, PDB ID: 1RY2, sequence identity: 44.73%). Secondary structures of *Sa*Acs1 are visually depicted as arrows (β-strands) and boxes (α-helices and 310/η-helices). 310/η-helices are normally considered as an extension of α-helices. Sequence motifs conserved among members of the AMP-forming superfamily are boxed and labeled A1 through A10. The yellow box indicates the residues where the beginning of the loop that interacts with the CoA molecule in the CoA binding pocket begins. Shading is based on the Blosum62 matrix with a threshold of 1, with residues that are 100% similar shaded black, those that are 80-100% similar are shaded charcoal, and those that are 60 to 80% similar are shaded gray.

**Table S1.** List of all AMP-forming Acs proteins deposited in the Protein Data Bank with the respective ligands and the conformation of the Acs structure.

| PDB ID | Organism Name | Ligands | Conformation | Ref |
|---|---|---|---|---|
| 1PG4 | *Salmonella enterica* (*Se*) | Propyl-AMP, CoA | Thioester | [1] |
| 2P2F | *Salmonella enterica* | Acetate, AMP, CoA | Thioester | [2] |
| 5JRH | *Salmonella enterica* | cAMP, CoA | Thioester | [3] |
| 5K85 | *Cryptococcus neoformans* (*Cn*) | CoA, propyl-AMP | Thioester | [4] |
| 5IFI | *Cryptococcus neoformans* | Propyl-AMP | Thioester | [4] |
| 5K8F | *Cryptococcus neoformans* | ATP | Adenylate | [4] |
| 7KNO | *Cryptococcus neoformans* | Ethyl-AMP | Adenylate | [4] |
| 7KNP | *Cryptococcus neoformans* | Butyl-AMP | Adenylate | [4] |
| 7L4G | *Cryptococcus neoformans* | Acetyl-AMP | Adenylate/Thioester | [4] |
| 5U29 | *Cryptococcus neoformans* | 5'-O-(acetylsulfamoyl)adenosine | Adenylate | [4] |
| 5VPV | *Cryptococcus neoformans* | none | Apo | [4] |
| 1RY2 | *Saccharomyces cerevisiae* (*Sc*) | AMP | Adenylate | [5] |
| 7MMZ | *Legionella pneumophila* (*Lp*) | Ethyl-AMP | Thioester | [6] |
| 7KQ6 | *Coccidioides immitis* (*Ci*) | Propyl-AMP | Thioester | n/a |
| 7L3Q | *Coccidioides immitis* | Methyl-AMP, CoA | Thioester | n/a |
| 7KVY | *Coccidioides immitis* | Ethyl-AMP, CoA | Thioester | n/a |
| 7KDN | *Aspergillus fumigatus* (*Af*) | Propyl-AMP | Thioester | n/a |
| 7KDS | *Candida albicans* (*Ca*) | Propyl-AMP | Thioester | n/a |
| 7KCP | *Coccidioides posadasii* (*Cp*) | Propyl-AMP | Thioester | n/a |
| 8SF3 | *Leishmania infantum* (*Li*) | Acetate, AMP, CoA | Thioester | [7] |
| 8U2R | *Leishmania infantum* | Ethyl-AMP | Thioester | [7] |
| 8U2S | *Leishmania infantum* | Ethyl-AMP | Thioester | [7] |
| 8U2T | *Leishmania infantum* | AMP, CoA | Thioester | [7] |
| 8U2U | *Leishmania infantum* | AMP, CoA | Thioester | [7] |

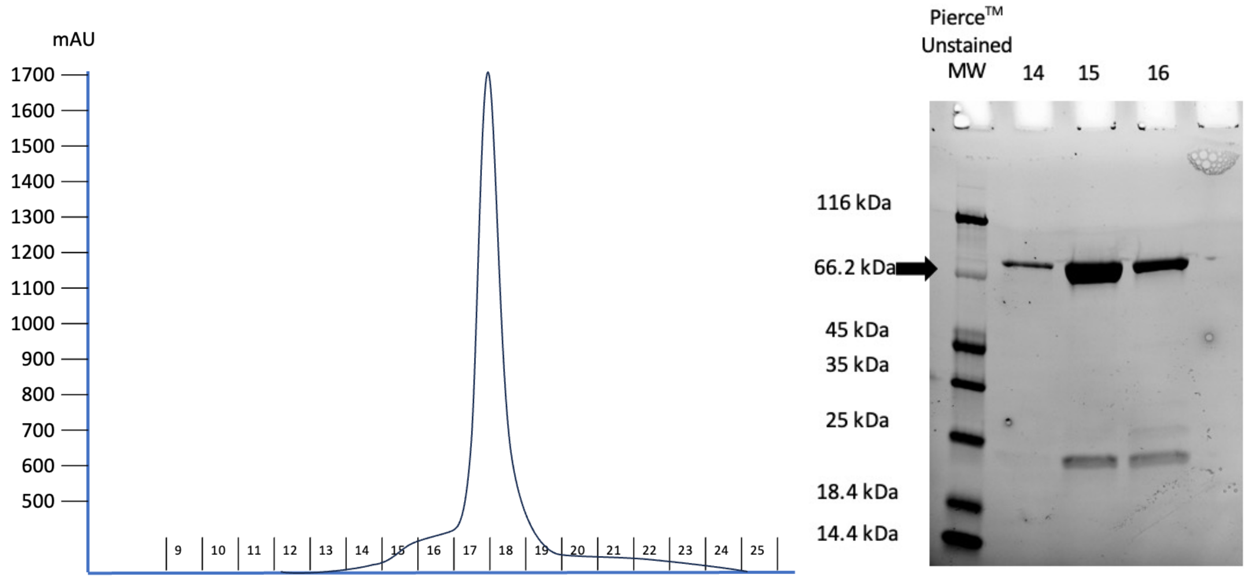


**Figure S5.** Overall purity of *Sa*Acs1 after nickel-affinity chromatography and size exclusion chromatography. Size-exclusion chromatography (SEC) chromatogram showing the UV absorption (mAU) on the y-axis and fraction numbers on the x-axis.  The first lane contains the molecular weight marker (Pierce™ Unstained Protein Molecular Weight Ladder) and lanes 2-4 contain SEC fractions which were pooled together. The black arrow indicates the molecular weight for *Sa*Acs1.

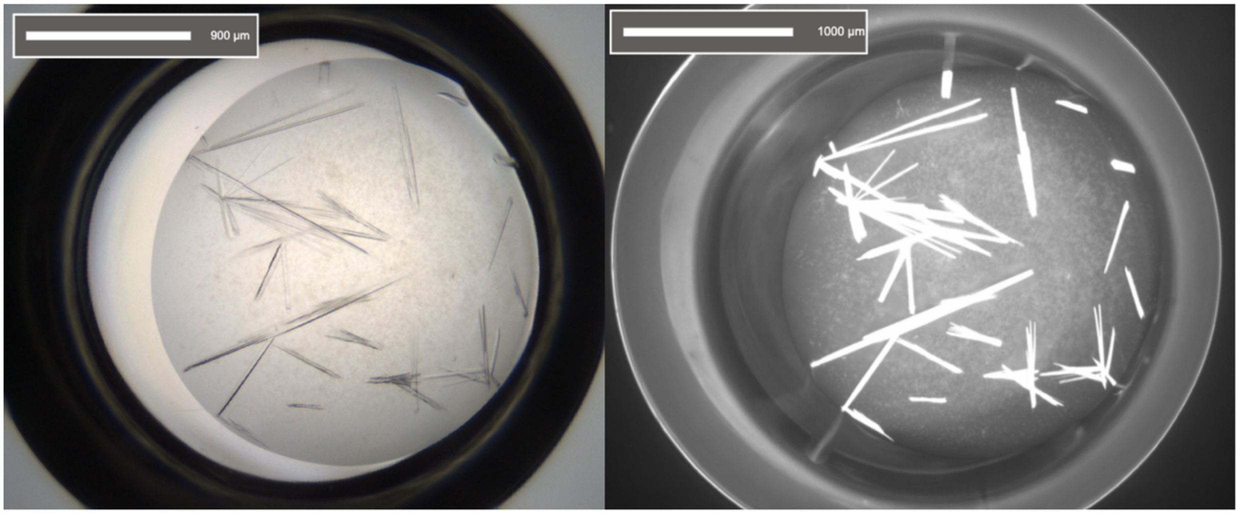


**Figure S6.** Visible light (left) and UV light (right) images of *Sa*Acs1$^{WT}$ crystals.

**Table S2.** Summary of crystal parameters, data collection, model, and refinement statistics for *Sa*Acs1$^{WT}$. Statistics for the highest-resolution shell are shown in parentheses.

| | ***Sa*Acs1$^{WT}$** | |
|---|---|---|
| **PDB ID** | 9Y8G | |
| **Resolution range** | 39.451 – 2.198 Å<br>(2.276 – 2.198 Å) | |
| **Space group** | C 2 2 $2_1$ | |
| **Unit cell** | **a:** 121.445 Å | **α:** 90° |
| | **b:** 175.979 Å | **β:** 90° |
| | **c:** 127.944 Å | **γ:** 90° |
| **Total Reflections** | 139166 (9459) | |
| **Unique Reflections** | 134417 (9235) | |
| **Completeness (%)** | 99.74 (97.10) | |
| **Mean I/sigma(I)** | 5.52 (1.55) | |
| **Wilson B-factor** | 29.23 | |
| **Reflections used in refinement** | 69674 (4795) | |
| **Reflections used for R-free** | 2000 (138) | |
| **R-work** | 0.1963 (0.2621) | |
| **R-free** | 0.2348 (0.3206) | |
| **Number of non-hydrogen atoms** | 10655 | |
| **macromolecules** | 10271 | |
| **ligands** | 53 | |
| **solvent** | 331 | |
| **Protein residues** | 1305 | |
| **RMS (bonds)** | 0.0007 | |
| **RMS (angles)** | 0.71 | |
| **Ramachandran favored (%)** | 97.08 | |
| **Ramachandran allowed (%)** | 2.77 | |
| **Ramachandran outliers (%)** | 0.15 | |
| **Rotamer outliers (%)** | 0.56 | |
| **Clashscore** | 2.44 | |
| **Average B-factor** | 31.58 | |
| **macromolecules** | 31.73 | |
| **ligands** | 20.95 | |
| **solvent** | 28.49 | |

**Table S3.** Summary of crystal parameters, data collection, model, and refinement statistics for *Sa*Acs1$^{G196E}$ and *Sa*Acs1$^{G196E/T197G}$. Statistics for the highest-resolution shell are shown in parentheses.

| | ***Sa*Acs1$^{G196E}$** | | ***Sa*Acs1$^{G196E/T197G}$** | |
|---|---|---|---|---|
| **PDB ID** | 36UO | | 36UT | |
| **Resolution range** | 39.39 – 1.699 Å<br>(1.74 – 1.7 Å) | | 29.74 – 2.301 Å<br>(2.36 – 2.3 Å) | |
| **Space group** | C 2 2 $2_1$ | | C 2 2 $2_1$ | |
| **Unit cell** | **a:** 121.266 Å | **α:** 90° | **a:** 120.530 Å | **α:** 90° |
| | **b:** 175.346 Å | **β:** 90° | **b:** 174.891 Å | **β:** 90° |
| | **c:** 128.423 Å | **γ:** 90° | **c:** 128.002 Å | **γ:** 90° |
| **Total Reflections** | 295003 (20207) | | 59936 (4262) | |
| **Unique Reflections** | 287346 (19853) | | 59887 (4193) | |
| **Completeness (%)** | 99.31 (97.06) | | 99.57 (98.17) | |
| **Mean I/sigma(I)** | 10.62 (1.60) | | 10.70 (1.34) | |
| **Wilson B-factor** | 25.01 | | 33.34 | |
| **Reflections used in refinement** | 148788 (10287) | | 59887 (4193) | |
| **Reflections used for R-free** | 2000 (138) | | 1997 (140) | |
| **R-work** | 0.2185 (0.3296) | | 0.2228 (0.2950) | |
| **R-free** | 0.2558 (0.3823) | | 0.2757 (0.3673) | |
| **Number of non-hydrogen atoms** | 10827 | | 10541 | |
| **macromolecules** | 10271 | | 10257 | |
| **ligands** | 53 | | 53 | |
| **solvent** | 503 | | 232 | |
| **Protein residues** | 1305 | | 1303 | |
| **RMS (bonds)** | 0.0013 | | 0.002 | |
| **RMS (angles)** | 1.17 | | 0.49 | |
| **Ramachandran favored (%)** | 96.85 | | 94.07 | |
| **Ramachandran allowed (%)** | 3.00 | | 5.54 | |
| **Ramachandran outliers (%)** | 0.15 | | 0.38 | |
| **Rotamer outliers (%)** | 0.93 | | 0.66 | |
| **Clashscore** | 3.09 | | 3.77 | |
| **Average B-factor** | 29.77 | | 36.86 | |
| **macromolecules** | 29.76 | | 37.00 | |
| **ligands** | 19.21 | | 26.14 | |
| **solvent** | 31.00 | | 33.02 | |

**Table S4.** Summary of crystal parameters, data collection, model, and refinement statistics for *Sa*Acs1$^{K202E}$ and *Sa*Acs1$^{D527P}$. Statistics for the highest-resolution shell are shown in parentheses.

| | ***Sa*Acs1$^{K202E}$** | | ***Sa*Acs1$^{D527P}$** | |
|---|---|---|---|---|
| **PDB ID** | 36UR | | 36US | |
| **Resolution range** | 39.62 – 2.098 Å (2.15 – 2.1 Å) | | 39.43 – 3.047 Å (3.12 – 3.05 Å) | |
| **Space group** | C 2 2 2$_1$ | | C 2 2 2$_1$ | |
| **Unit cell** | **a:** 121.822 Å | **α:** 90° | **a:** 121.375 Å | **α:** 90° |
| | **b:** 176.292 Å | **β:** 90° | **b:** 175.565 Å | **β:** 90° |
| | **c:** 129.400 Å | **γ:** 90° | **c:** 126.826 Å | **γ:** 90° |
| **Total Reflections** | 159334 (10947) | | 51128 (3452) | |
| **Unique Reflections** | 154406 (10717) | | 48805 (3349) | |
| **Completeness (%)** | 98.21 (96.04) | | 98.71 (93.22) | |
| **Mean I/sigma(I)** | 7.42 (1.05) | | 3.52 (0.81) | |
| **Wilson B-factor** | 36.40 | | 63.64 | |
| **Reflections used in refinement** | 79925 (5509) | | 25906 (1720) | |
| **Reflections used for R-free** | 2000 (138) | | 1998 (133) | |
| **R-work** | 0.2250 (0.3002) | | 0.2166 (0.3605) | |
| **R-free** | 0.2771 (0.3575) | | 0.2597 (0.3849) | |
| **Number of non-hydrogen atoms** | 10324 | | 10326 | |
| **macromolecules** | 10271 | | 10269 | |
| **ligands** | 53 | | 53 | |
| **solvent** | 210 | | 4 | |
| **Protein residues** | 1305 | | 1305 | |
| **RMS (bonds)** | 0.007 | | 0.002 | |
| **RMS (angles)** | 0.86 | | 0.43 | |
| **Ramachandran favored (%)** | 96.16 | | 95.62 | |
| **Ramachandran allowed (%)** | 3.69 | | 4.30 | |
| **Ramachandran outliers (%)** | 0.15 | | 0.08 | |
| **Rotamer outliers (%)** | 0.74 | | 0.00 | |
| **Clashscore** | 3.48 | | 3.09 | |
| **Average B-factor** | 37.35 | | 60.09 | |
| **macromolecules** | 37.42 | | 60.17 | |
| **ligands** | 25.07 | | 46.13 | |
| **solvent** | 35.39 | | 29.84 | |

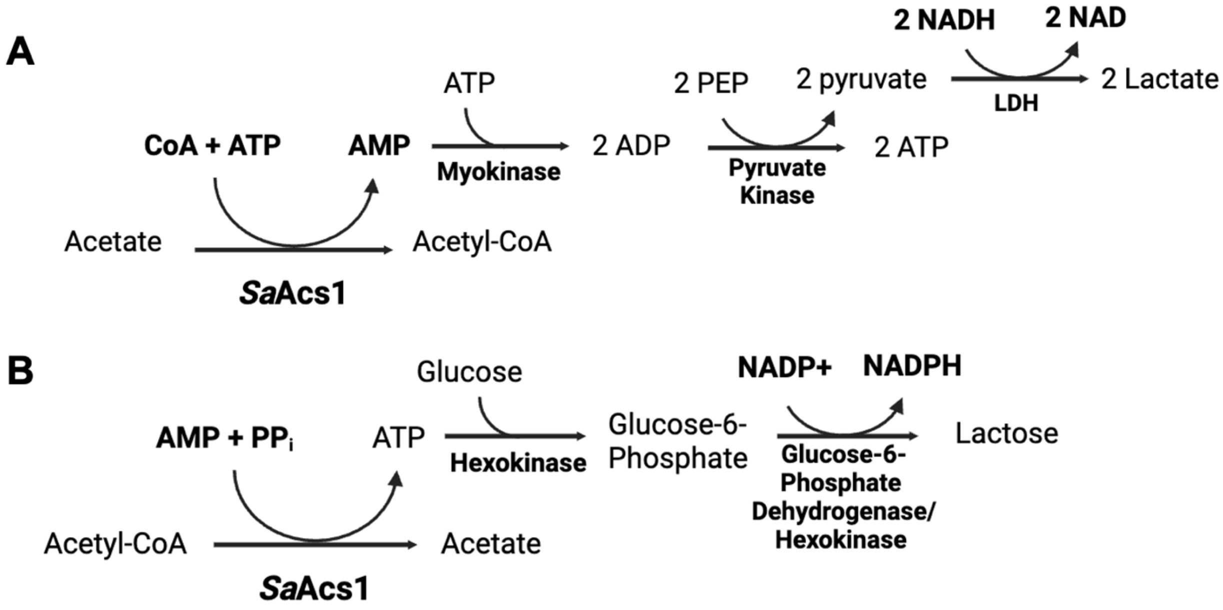


**Figure S7.** The *Sa*Acs1 (**A**) AMP-forming reaction and (**B**) ATP-forming reaction for activity and kinetics assays.

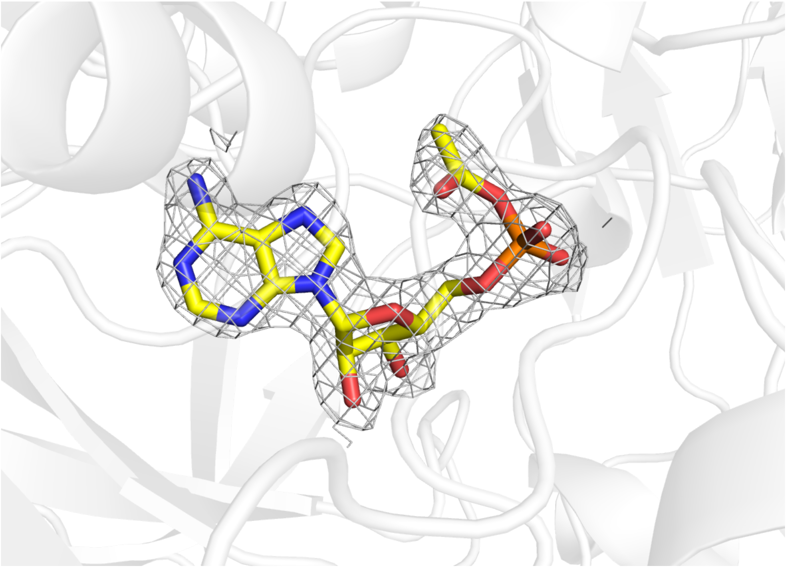

**Figure S8.** Active site of *Sa*Acs1$^{WT}$ with acetyl-AMP shown in yellow. Electron density for acetyl-AMP is shown at 1σ of a $2F_O$-$F_C$ map.

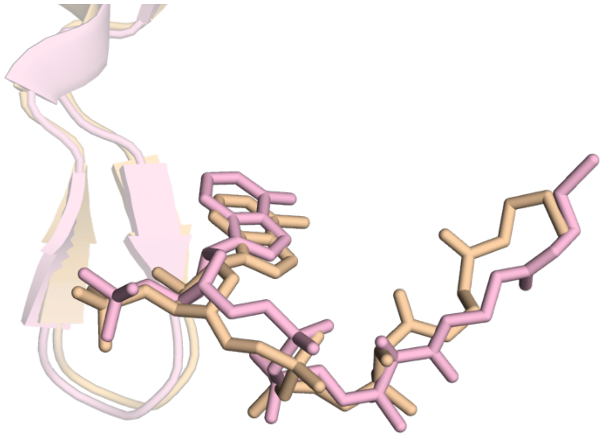

**Figure S9.** View of the CoA loop residing in the CoA pocket that interacts with the nucleotide end of the CoA molecule. The structures of *Se*Acs (salmon, PDB ID: 2P2F, chain A) and *Cn*Acs (tan, PDB ID: 5K85, chain C) along with the respective CoA molecules from the *Se*Acs and *Cn*Acs structures.

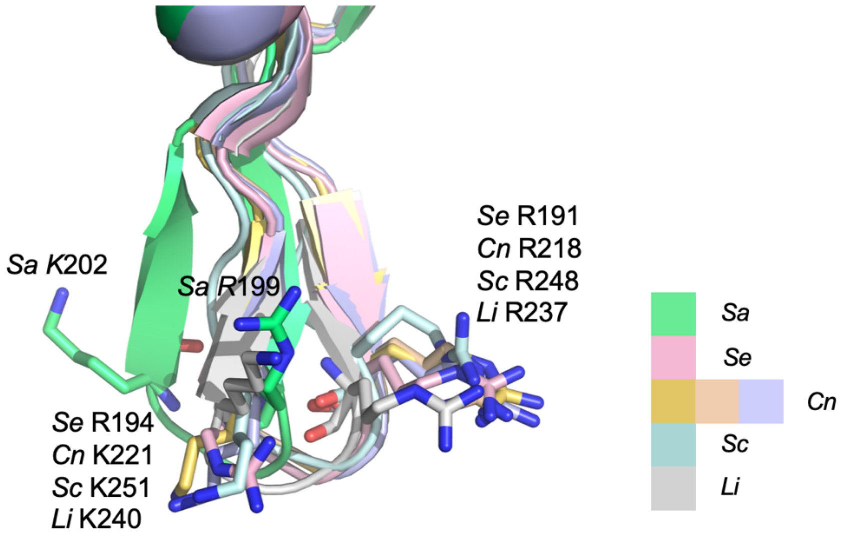


**Figure S10.** View of the CoA-binding loop alignment shown with the conserved residues in the loop of each respective structure. Alignment of multiple Acs structures resolved in the apo, adenylation (AD), thioesterification (TE) conformations with various bound ligands. Structures of *Se*Acs (salmon, PDB ID 2P2F chain A, TE – bound CoA), *Cn*Acs (light orange, PDB ID 7L4G, chain C, TE – without CoA), *Cn*Acs (tan, PDB ID 5K85, chain C, TE - bound CoA), *Cn*Acs (lavender, PDB ID 5VPV, chain A, apo), *Sc*Acs (light cyan, PDB ID 1RY2, chain A, AD), *Li*Acs (light gray, PDB ID 8U2R, chain A, TE – without CoA) compared to *Sa*Acs1 (lime green, chain A, AD). Legend includes the color representation for each structure.

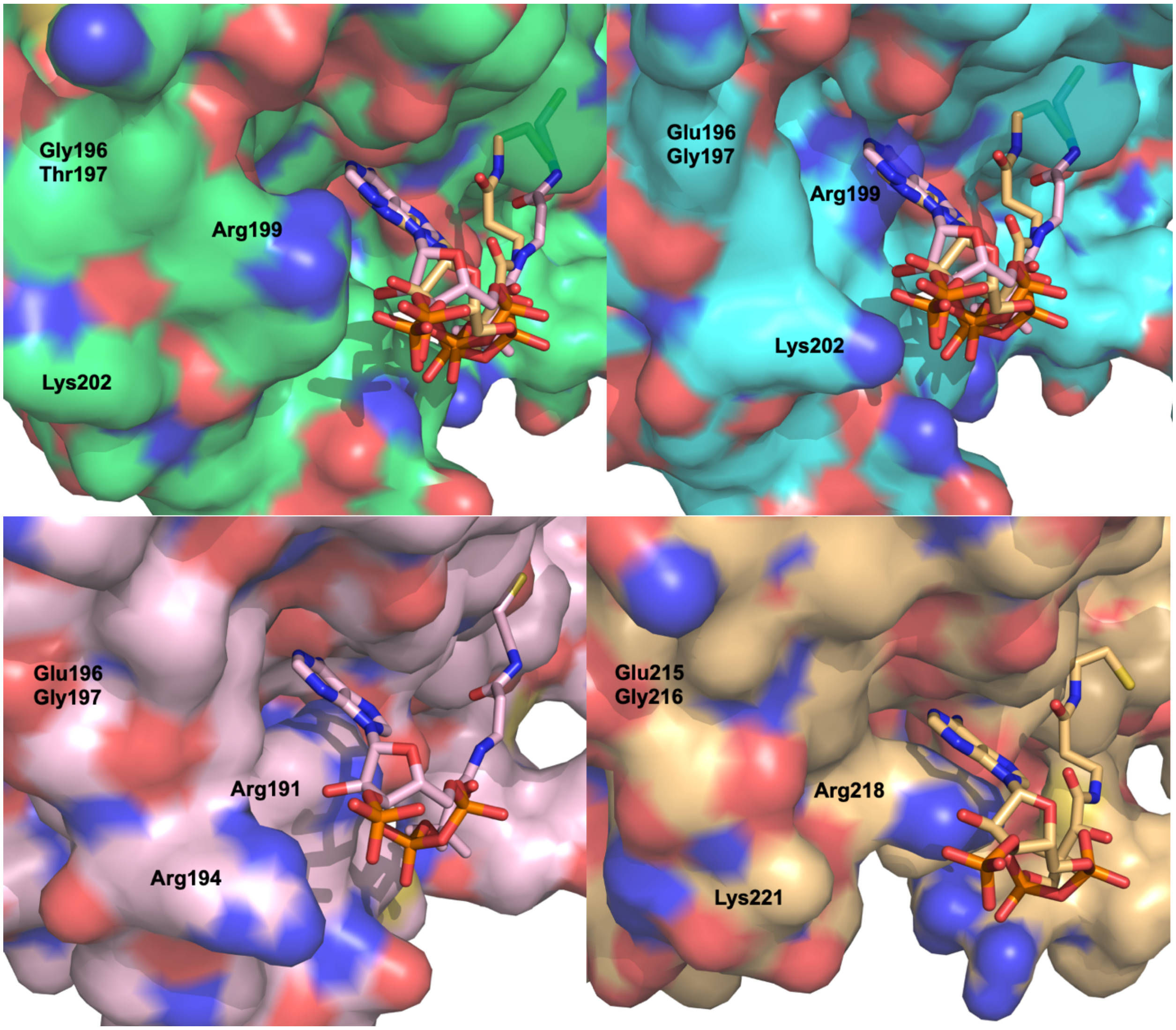


**Figure S11.** Comparison of the CoA-binding pocket surface with respective residues within the CoA loop for *Sa*Acs1 (lime green, chain A), *Se*Acs (salmon, PDB ID: 2P2F, chain A), and *Cn*Acs (tan, PDB ID: 5K8F, chain C). The *Sa*Acs1 structure includes overlays of CoA from *Se*Acs (salmon, PDB ID: 2P2F, chain A) and *Cn*Acs (tan, PDB ID: 5K8F, chain C). The nitrogen and oxygen atoms for all residues and ligands are shown in blue and red, respectively.

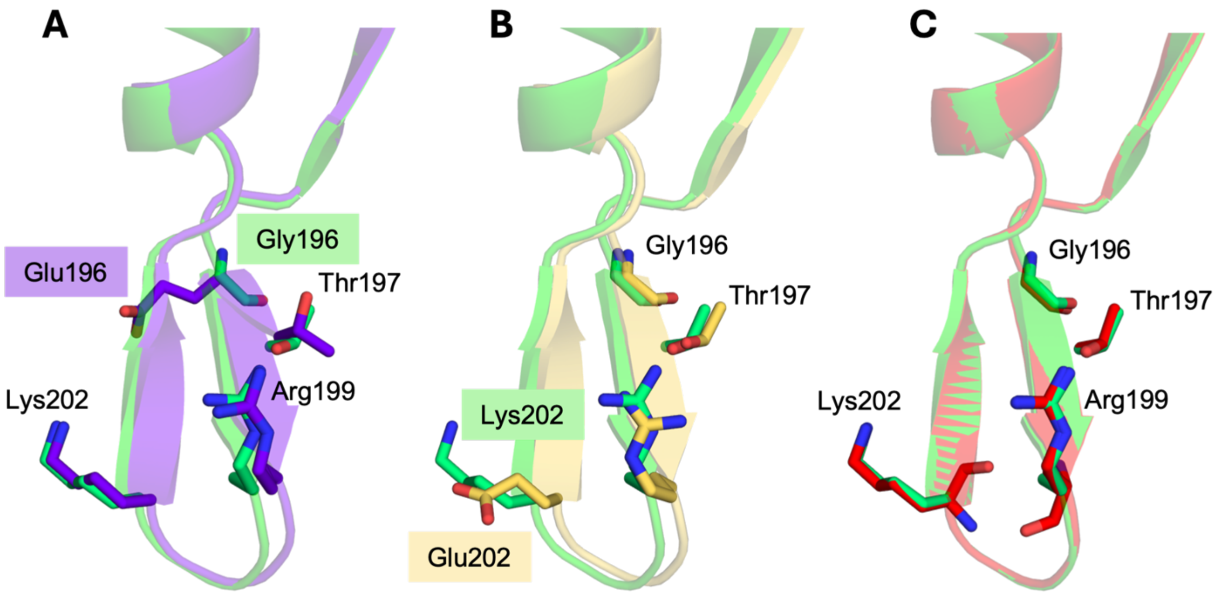


**Figure 12.** Alignment of the CoA-binding loops from *Sa*Acs1 variants (**A**) G196E (purple) (**B**) K202E (yellow) (**C**) D527P (red).

### Kinetic Parameter Determination for Acs Enzymes

#### Apparent Steady-State Kinetic Parameters and Catalytic Bias Analysis

Acetyl-CoA synthetase (Acs) catalyzes a reversible ordered Bi-Uni-Uni-Bi ping-pong reaction through two chemically distinct half-reactions: formation of an enzyme-bound acetyl-adenylate intermediate followed by CoA-dependent thioester formation (**Figures S2 and S3**).[8] Unlike a classical single-substrate Michaelis-Menten enzyme, the observed steady-state kinetic parameters for Acs depend on multiple substrate-binding, conformational, catalytic, and product-release steps. Consequently, the experimentally determined $K_M$ and $k_{cat}$ values do not correspond to individual microscopic binding or catalytic constants, but instead represent **apparent steady-state parameters** that describe the behavior of the complete catalytic cycle under a defined set of assay conditions. This distinction is well-established for multi-substrate enzyme mechanisms.[9-11]

In this study, kinetic parameters were determined by varying a single substrate while maintaining the remaining substrates at saturating concentrations, following the experimental strategy commonly employed for Acs enzymes and other members of the adenylate-forming enzyme superfamily. Under these conditions, the measured $K_M$, $k_{cat}$, and $k_{cat}/K_M$ values should be interpreted as operational or apparent kinetic parameters. Their numerical values therefore depend not only on the intrinsic microscopic rate constants of the catalytic mechanism but also on the concentrations chosen for the remaining substrates, the degree of co-substrate saturation achieved experimentally, and the step that limits turnover under those assay conditions.

Importantly, this distinction does not diminish the utility of these measurements. Although apparent kinetic parameters cannot generally be interpreted as microscopic binding affinities or elementary catalytic rate constants, they provide a consistent and experimentally reproducible description of overall enzyme performance under identical assay conditions. For comparative studies such as the present work, where all enzymes and variants were characterized using the same experimental protocol, differences in the apparent kinetic parameters directly report how mutations redistribute catalytic performance within the catalytic cycle.

#### Apparent Haldane Relationship

For a reversible enzyme, the classical Haldane relationship relates the equilibrium constant to the kinetic constants of the complete microscopic mechanism. In multi-substrate reactions, however, the exact Haldane relationship depends not only on Michaelis constants but also on inhibition constants and other kinetic parameters describing substrate and product binding. For ordered and ping-pong mechanisms these additional terms arise naturally from the steady-state rate equation and generally cannot be determined from single-substrate Michaelis-Menten experiments alone.

If the Michaelis constants are substituted for the corresponding microscopic constants, an **apparent Haldane ratio** may be written as

$$K_{\mathrm{app}} = \frac{k_{\mathrm{cat},f}\, K_M^{\mathrm{PP_i}}\, K_M^{\mathrm{AMP}}\, K_M^{\mathrm{Ac-CoA}}}{k_{\mathrm{cat},r}\, K_M^{\mathrm{acetate}}\, K_M^{\mathrm{ATP}}\, K_M^{\mathrm{CoA}}}.$$

This expression is not intended to estimate the thermodynamic equilibrium constant of the reaction. Instead, it provides a convenient composite kinetic parameter that summarizes the experimentally observed distribution of apparent catalytic efficiencies between the ATP-forming and AMP-forming directions.

Because a formal $k_{cat}$ is defined only once for each reaction direction, the individual $k_{cat}$ values obtained by varying different substrates should be viewed as independent experimental estimates of the same apparent turnover number rather than distinct microscopic catalytic constants. The CoA- and acetyl-CoA-dependent measurements were used to represent the forward and reverse turnover numbers because these substrates participate directly in the thioester-forming half-reaction, which is thought to limit overall turnover for the forward reaction and, by microscopic reversibility, is expected to contribute similarly in the reverse direction. We emphasize, however, that alternative choices produce only modest quantitative differences and do not alter the qualitative comparison among enzymes.

**Catalytic Bias**

Because the primary objective of this work is to compare the relative specialization of homologous Acs enzymes rather than determine a complete microscopic kinetic mechanism, we use a effective catalytic bias index, $\beta$, based directly on the measured catalytic efficiencies:

$$\beta = \log_{10}\left[\frac{\left(\frac{k_{\text{cat}}}{K_M}\right)_{\text{ATP}}\left(\frac{k_{\text{cat}}}{K_M}\right)_{\text{acetate}}\left(\frac{k_{\text{cat}}}{K_M}\right)_{\text{CoA}}}{\left(\frac{k_{\text{cat}}}{K_M}\right)_{\text{AMP}}\left(\frac{k_{\text{cat}}}{K_M}\right)_{\text{PP}_\text{i}}\left(\frac{k_{\text{cat}}}{K_M}\right)_{\text{Ac-CoA}}}\right].$$

Positive values of $\beta$ indicate a kinetic bias toward the canonical AMP-forming direction, whereas negative values indicate bias toward the ATP-forming direction. Because this metric is calculated directly from the experimentally measured catalytic efficiencies, it avoids assumptions regarding the microscopic mechanism while providing a quantitative description of how catalytic efficiency is redistributed between the two directions of the reversible reaction.

**Table S5.** The $K_M$ (μM), $k_{cat}$ ($s^{-1}$), and $k_{cat}/K_m$ ($\mu M^{-1}s^{-1}$) with the symmetrical confidence interval (standard error) for each substrate in both reactions for *Sa*Acs1$^{WT}$, *Sa*Acs1 CoA-binding loop variants, *Se*Acs$^{WT}$, and *Cn*Acs$^{WT}$. ND=not detectible.

| | $K_M$ (μM) | | | | | |
|---|---|---|---|---|---|---|
| | Acetate | ATP | CoA | PPi | AMP | Acetyl-CoA |
| *Sa*Acs1$^{WT}$ | 73.9 ± 18.54 | 303.6 ± 48.37 | 61.94 ± 24.11 | 99.65 ± 19.89 | 102.5 ± 27.33 | 109.3 ± 20.23 |
| *Sa*Acs1$^{G196E}$ | 47.22 ± 12.48 | 427 ± 68.89 | 51.05 ± 11.84 | 105.6 ± 27.26 | 149.6 ± 21.39 | 115.2 ± 18.22 |
| *Sa*Acs1$^{G196E/T197G}$ | 81.32 ± 15.77 | 136.6 ± 30.50 | 25.61 ± 7.51 | 106.5 ± 16.07 | 115.7 ± 26.52 | 150.3 ± 20.02 |
| *Sa*Acs1$^{G196E/T197G/K202R}$ | 89.09 ± 22.78 | 98.62 ± 26.06 | 24.38 ± 6.25 | 52.12 ± 6.55 | 26.06 ± 7.15 | 31.36 ± 3.72 |
| *Se*Acs$^{WT}$ | 96.5 ± 17.56 | 93.5 ± 20.73 | 40.9 ± 4.34 | 143.5 ± 43.47 | 176.6 ± 51.81 | 181.2 ± 36.08 |
| *Se*Acs$^{E188G/G189T}$ | 75.46 ± 16.26 | 106.1 ± 24.41 | 36.66 ± 10.2 | ND | ND | ND |
| *Se*Acs$^{E188G/G189T/R194K}$ | 42.69 ± 14.82 | 108 ± 28.74 | 28.28 ± 10.28 | ND | ND | ND |
| *Cn*Acs$^{WT}$ | 118.6 ± 34.27 | 139.8 ± 24.58 | 142.7 ± 31.44 | 252.4 ± 87.14 | 171.9 ± 42.22 | 177.2 ± 15.21 |

| | $k_{cat}$ ($s^{-1}$) | | | | | |
|---|---|---|---|---|---|---|
| | Acetate | ATP | CoA | PPi | AMP | Acetyl-CoA |
| *Sa*Acs1$^{WT}$ | 97.25 ± 4.52 | 121.7 ± 3.59 | 133.7 ± 3.54 | 115.1 ± 6.09 | 101.6 ± 4.91 | 140.1 ± 8.29 |
| *Sa*Acs1$^{G196E}$ | 120.6 ± 5.32 | 106.1 ± 3.56 | 132.4 ± 7.44 | 88.79 ± 6.24 | 91.04 ± 2.63 | 109.6 ± 5.67 |
| *Sa*Acs1$^{G196E/T197G}$ | 116.8 ± 4.15 | 117.4 ± 5.32 | 118.8 ± 6.35 | 83.07 ± 3.53 | 72.66 ± 3.28 | 147.4 ± 7.37 |
| *Sa*Acs1$^{G196E/T197G/K202R}$ | 105.8 ± 5.05 | 102 ± 4.26 | 117.2 ± 5.4 | 43.89 ± 1.07 | 40.98 ± 1.22 | 44.15 ± 1.99 |
| *Se*Acs$^{WT}$ | 83.86 ± 1.9 | 86.08 ± 3.76 | 119 ± 2.63 | 73.22 ± 5.05 | 74.45 ± 4.84 | 92.58 ± 7.42 |
| *Se*Acs$^{E188G/G189T}$ | 82.34 ± 3.21 | 81.41 ± 3.42 | 82.38 ± 4.62 | ND | ND | ND |
| *Se*Acs$^{E188G/G189T/R194K}$ | 134.7 ± 7.49 | 141.1 ± 6.89 | 141.3 ± 9.62 | ND | ND | ND |
| *Cn*Acs$^{WT}$ | 146.1 ± 5.73 | 105.4 ± 3.46 | 210.7 ± 15.88 | 136.6 ± 14.81 | 116.6 ± 7.04 | 130 ± 7.07 |

| | $k_{cat}/K_M$ ($\mu M^{-1}s^{-1}$) | | | | | |
|---|---|---|---|---|---|---|
| | Acetate | ATP | CoA | PPi | AMP | Acetyl-CoA |
| *Sa*Acs1$^{WT}$ | 1.32 ± 0.24 | 0.4 ± 0.07 | 2.16 ± 0.15 | 1.15 ± 0.31 | 0.99 ± 0.18 | 1.28 ± 0.41 |
| *Sa*Acs1$^{G196E}$ | 2.55 ± 0.42 | 0.24 ± 0.05 | 2.59 ± 0.63 | 0.84 ± 0.23 | 0.61 ± 0.12 | 0.95 ± 0.31 |
| *Sa*Acs1$^{G196E/T197G}$ | 1.44 ± 0.26 | 0.86 ± 0.17 | 4.64 ± 0.85 | 0.78 ± 0.22 | 0.63 ± 0.12 | 0.98 ± 0.37 |
| *Sa*Acs1$^{G196E/T197G/K202R}$ | 1.19 ± 0.22 | 1.03 ± 0.19 | 4.81 ± 0.86 | 0.84 ± 0.16 | 1.57 ± 0.17 | 1.41 ± 0.53 |
| *Se*Acs$^{WT}$ | 0.9 ± 0.11 | 0.92 ± 0.18 | 2.91 ± 0.61 | 0.51 ± 0.21 | 0.42 ± 0.09 | 0.51 ± 0.21 |
| *Se*Acs$^{E188G/G189T}$ | 1.09 ± 0.20 | 0.77 ± 0.14 | 2.25 ± 0.45 | ND | ND | ND |
| *Se*Acs$^{E188G/G189T/R194K}$ | 3.16 ± 0.51 | 1.31 ± 0.24 | 5 ± 0.94 | ND | ND | ND |
| *Cn*Acs$^{WT}$ | 1.23 ± 0.17 | 0.75 ± 0.14 | 1.48 ± 0.51 | 0.54 ± 0.17 | 0.68 ± 0.17 | 0.73 ± 0.46 |

**Table S6.** The $K_M$ (µM), $k_{cat}$ ($s^{-1}$), and $k_{cat}/K_m$ ($\mu M^{-1}s^{-1}$) with the symmetrical confidence interval (standard error) for each substrate in both reactions for *Sa*Acs1$^{WT}$ compared to *Sa*Acs1 CoA loop variants. ND=not detectible.

| | $K_M$ (µM) | | | | | |
|---|---|---|---|---|---|---|
| | Acetate | ATP | CoA | PPi | AMP | Acetyl-CoA |
| *Sa*Acs1$^{R199A}$ | 74.21 ± 27.31 | 346.9 ± 64.91 | 43.34 ± 11.72 | ND | ND | ND |
| *Sa*Acs1$^{R199E}$ | 42.41 ± 7.47 | 254.9 ± 77.42 | 27.12 ± 5.63 | ND | ND | ND |
| *Sa*Acs1$^{K202A}$ | 80.08 ± 16.32 | 349 ± 102.4 | 130.3 ± 44.21 | ND | ND | ND |
| *Sa*Acs1$^{K202R}$ | 142.4 ± 38.39 | 106 ± 28.03 | 22.17 ± 7.14 | 150.3 ± 18.88 | 145.8 ± 19.26 | 195.1 ± 24.08 |
| *Sa*Acs1$^{K202E}$ | 21.54 ± 3.53 | 331.6 ± 114.3 | 58.16 ± 14.61 | ND | ND | ND |
| *Sa*Acs1$^{K202S}$ | 163.6 ± 48.14 | 424.7 ± 89.89 | 116.2 ± 10.74 | 110.4 ± 17.97 | 169± 22.78 | 157.2 ± 32.63 |
| *Sa*Acs1$^{K202Q}$ | 72.13 ± 14.03 | 362.6 ± 140 | 85.83 ± 11.64 | 98.31 ± 6.67 | 108.6 ± 19.43 | 113.2 ± 30.05 |
| *Sa*Acs1$^{WT}$ | 73.9 ± 18.54 | 303.6 ± 48.37 | 61.94 ± 24.11 | 99.65 ± 19.89 | 102.5 ± 27.33 | 109.3 ± 20.23 |

| | $k_{cat}$ ($s^{-1}$) | | | | | |
|---|---|---|---|---|---|---|
| | Acetate | ATP | CoA | PPi | AMP | Acetyl-CoA |
| *Sa*Acs1$^{R199A}$ | 125.5 ± 8.54 | 133.5 ± 4.53 | 149.5 ± 9.26 | ND | ND | ND |
| *Sa*Acs1$^{R199E}$ | 80.81 ± 2.3 | 82.6 ± 4.39 | 90.79 ± 3.72 | ND | ND | ND |
| *Sa*Acs1$^{K202A}$ | 75.3 ± 2.85 | 76.85 ± 4.37 | 96.31 ± 10.52 | ND | ND | ND |
| *Sa*Acs1$^{K202R}$ | 100.6 ± 7.06 | 90.35 ± 4.85 | 98.79 ± 5.78 | 92.86 ± 3.69 | 76.95 ± 2.04 | 102.1 ± 5.2 |
| *Sa*Acs1$^{K202E}$ | 70.02 ± 1.48 | 67.31 ±4.42 | 85.8 ± 5.8 | ND | ND | ND |
| *Sa*Acs1$^{K202S}$ | 118.2 ± 7.34 | 138.7 ± 5.73 | 167.7 ± 4.45 | 117.1 ± 5.29 | 126.6 ± 3.58 | 153.2 ± 11.96 |
| *Sa*Acs1$^{K202Q}$ | 131.3 ± 4.65 | 129.2 ± 9.05 | 169.8 ± 5.63 | 140.3 ± 2.51 | 120.5 ± 3.97 | 139.4 ± 12.02 |
| *Sa*Acs1$^{WT}$ | 97.25 ± 4.52 | 121.7 ± 3.59 | 133.7 ± 3.54 | 115.1 ± 6.09 | 101.6 ± 4.91 | 140.1 ± 8.29 |

| | $k_{cat}/K_M$ ($\mu M^{-1}s^{-1}$) | | | | | |
|---|---|---|---|---|---|---|
| | Acetate | ATP | CoA | PPi | AMP | Acetyl-CoA |
| *Sa*Acs1$^{R199A}$ | 1.69 ± 0.31 | 0.38 ± 0.07 | 3.45 ± 0.779 | ND | ND | ND |
| *Sa*Acs1$^{R199E}$ | 1.91 ± 0.31 | 0.32 ± 0.06 | 3.35 ± 0.66 | ND | ND | ND |
| *Sa*Acs1$^{K202A}$ | 0.94 ± 0.17 | 0.22 ± 0.04 | 0.74 ± 0.23 | ND | ND | ND |
| *Sa*Acs1$^{K202R}$ | 0.71 ± 0.18 | 0.85 ± 0.15 | 4.45 ± 0.81 | 0.62 ± 0.2 | 0.53 ± 0.11 | 0.52 ± 0.22 |
| *Sa*Acs1$^{K202E}$ | 3.25 ± 0.42 | 0.2 ± 0.04 | 1.48 ± 0.4 | ND | ND | ND |
| *Sa*Acs1$^{K202S}$ | 0.72 ± 0.15 | 0.33 ± 0.06 | 1.44 ± 0.41 | 1.06 ± 0.29 | 0.75 ± 0.16 | 0.97 ± 0.36 |
| *Sa*Acs1$^{K202Q}$ | 1.82 ± 0.33 | 0.35 ± 0.06 | 1.98 ± 0.37 | 1.41 ± 0.37 | 1.11 ± 0.2 | 1.23 ± 0.4 |
| *Sa*Acs1$^{WT}$ | 1.32 ± 0.24 | 0.4 ± 0.07 | 2.16 ± 0.15 | 1.15 ± 0.31 | 0.99 ± 0.18 | 1.28 ± 0.41 |
